# Interactions between a mechanosensitive channel and cell wall integrity signaling influence pollen germination in *Arabidopsis thaliana*

**DOI:** 10.1101/2021.08.24.457556

**Authors:** Yanbing Wang, Joshua Coomey, Kari Miller, Gregory S. Jensen, Elizabeth S. Haswell

## Abstract

Cells employ multiple systems to maintain cellular integrity, including mechanosensitive (MS) ion channels and the cell wall integrity (CWI) pathway. Here, we use pollen as a model system to ask how these different mechanisms are interconnected at the cellular level. MscS-Like (MSL)8 is an MS channel required to protect *Arabidopsis thaliana* pollen from osmotic challenges during *in vitro* rehydration, germination and tube growth. New CRISPR/Cas9 and artificial microRNA-generated *msl8* alleles produced unexpected pollen phenotypes, including the ability to germinate a tube after bursting, dramatic defects in cell wall structure and disorganized callose deposition at the germination site. We document complex genetic interactions between *MSL8* and two previously established components of the CWI pathway, *MARIS*, and *ANXUR1/2*. Overexpression of *MARIS^R240C^-FP* suppressed the bursting, germination, and callose deposition phenotypes of *msl8* mutant pollen. Null *msl8* alleles suppressed the internalized callose structures observed in *MARIS^R240C^-FP* lines. Similarly, *MSL8-YFP* overexpression suppressed bursting in the *anxur1/2* mutant background, while *anxur1/2* alleles reduced the strong rings of callose around ungerminated pollen grains in *MSL8-YFP* over-expressors. These data show that MS ion channels modulate callose deposition in pollen and provides evidence that cell wall and membrane surveillance systems coordinate in a complex manner to maintain cell integrity.

## INTRODUCTION

Cell mechanobiology includes the study of mechanical properties of cell constituents and the forces applied to or generated by cells and their microenvironment (Bidhendi & Geitmann 2019). Plant cells perceive and respond to multiple external or internal forces important for their growth, survival, and development (Monshausen & Haswell 2013; Trinh *et al*. 2021). Pollen, the male gamete of flowering plants, is an excellent model system for the study of plant cell mechanobiology due to its relatively simple structure, its unique biology, and the availability of *in vitro* growth systems (Johnson, Harper & Palanivelu 2019).This single cell harbors two sperm cells and germinates a tube to deliver the sperm to the female ovule for double fertilization, navigating multiple osmotic and mechanical challenges during rehydration, germination, tube growth, and fertilization (Firon, Nepi & Pacini 2012; Adhikari, Liu & Kasahara 2020).

The plasma membrane is an important platform for mechanoresponses and it is expected that changes in the physical properties of the membrane result from changes in turgor or in cell wall integrity (Ackermann & Stanislas 2020). A conserved mechanism for force perception at the membrane is the activation of mechanosensitive (MS) channels through increased membrane tension (Naismith & Booth 2012; Peyronnet, Tran, Girault & Frachisse 2014; Ranade, Syeda & Patapoutian 2015). In plants, members of the MscS-like (MSL) family of mechanosensitive ion channels are localized to mitochondrial, plastid, and plasma membranes (Basu & Haswell 2017), where they function as tension-regulated osmotic safety valves (Wilson, Jensen & Haswell 2011; Hamilton *et al*. 2015; Hamilton & Haswell 2017), some with additional complex functions (Veley *et al*. 2014; Maksaev, Shoots, Ohri & Haswell 2018). MSL8 is localized to the plasma membrane of pollen grains and tubes, and mediates mechanosensitive channel activities with a slight preference for anions (Hamilton *et al*. 2015). MSL8’s tension-sensitive ion transport activity is required to protect Arabidopsis pollen from osmotic challenges during *in vitro* pollen rehydration, germination and tube growth (Hamilton & Haswell 2017). Loss of MSL8 causes pollen grain to burst (Hamilton *et al*. 2015).

The plant cell wall provides tensile strength and protection against both internal and external mechanical stresses (Höfte & Voxeur 2017). The cell wall in a mature pollen surface comprises an inner intine wall, an outer exine wall, and a pollen coat. Callose, a polysaccharide in the form of β-1,3-glucan with some β-1,6-branches (Chen & Kim 2009), is abundant in immature pollen grains, but disappears during pollen maturation and desiccation (Xu, Zhang, Zhou & Yang 2016). Once a pollen grain has rehydrated and is about to germinate, tube emergence begins with the formation of a germination plaque (Johnson & McCormick 2001; Hoedemaekers *et al*. 2015). This lens-shaped plaque is typically restricted to one site on the grain periphery(Johnson & McCormick 2001; Hoedemaekers *et al*. 2015). It contains callose, which is thought to add flexibility to the cell wall and increase its load-bearing ability (Parre & Geitmann 2005), During germination, the germination plaque protrudes and extends, eventually forming the tip-growing tube that will eventually deliver the male gamete to the female egg cell (Hoedemaekers *et al*. 2015). A tightly controlled balance between cell wall deposition and internal pressure allows the pollen tube to rapidly reach the ovules for fertilization (Cameron & Geitmann 2018).

In plants, the cell wall integrity (CWI) signaling pathway serves to maintain mechanical homeostasis during cell expansion and in response to cell wall weakening (Gigli-Bisceglia, Engelsdorf & Hamann 2019). Members of the *Catharanthus roseus* RLK1–like (CrRLK1L) protein family are proposed to sense wall conditions and transduce information to cytosolic signaling factors (Li, Wu & Cheung 2016).The CrRLK1Ls involved CWI signaling pathway is vital to maintaining cellular integrity in multiple processes, including pollen grain germination and pollen tube growth (Li *et al*. 2016; Li & Yang 2018; Ge, Cheung & Qu 2019; Vogler, Santos-Fernandez, Mecchia & Grossniklaus 2019). In *Arabidopsis thaliana* pollen, ANXUR1 and ANXUR2 (ANX1/2) and Buddha’s Paper Seal (BUPS)1/2 are membrane localized CrRLK1L receptors. The ANX1/2-BUPS1/2 complex, as well as wall-anchored Leucine-rich repeat extensins (LRX8/9/11), bind the autocrine small peptides known as Rapid Alkalinization Factor (RALF)4/19 to sense alterations in the cell wall. These signals are transduced to downstream signaling components, including the receptor-like cytoplasmic kinase MARIS (MRI), to regulate pollen tube growth rate and maintain cell integrity (Liao *et al*. 2016; Li & Yang 2018; Ge *et al*. 2019). Mutants that disrupt the pathway, such as *anx1/2, mri, lrx8/9/11, bups1/2* and *ralf4/19*, show increased grain bursting (Boisson-Dernier *et al*. 2009, 2013; Boisson-Dernier, Franck, Lituiev & Grossniklaus 2015; Miyazaki *et al*. 2009; Liao *et al*. 2016; Ge *et al*. 2017; Fabrice *et al*. 2018; Sede *et al*. 2018) and some have elevated callose deposition (Fabrice *et al*. 2018; Sede *et al*. 2018). Over-expressing either *ANX1/2* or a dominant *MRI* allele (*MRI*^*R240C*^) results in membrane invagination and hyperaccumulation of cell wall material (Boisson-Dernier *et al*. 2013, 2015). MRI functions downstream of ANX1/2, as overexpression of MRI^R240C^ suppresses grain bursting in *anx1/2* mutants (Boisson-Dernier *et al*. 2015).

It has been proposed that the CWI pathway could act in synergy with mechanosensitive channels, to integrate mechanical information from the cell wall and plasma membrane and maintain cellular mechanical homeostasis (Hamant & Haswell 2017). The similar pollen bursting phenotypes of *msl8* mutants and CWI mutants supports this idea. Here we further test this hypothesis with new mutant alleles of *MSL8* and two components of the pollen CWI pathway. Our data indicate a novel function of *MSL8* in cell wall organization during pollen germination and suggest a complex connection between *MSL8* and pollen CWI in maintaining cell integrity during germination.

## RESULTS

### Creation of new *msl8* and *msl7 msl8* mutant lines

We previously characterized pollen hydration and germination defects in two *MSL8* T-DNA insertion mutants (Hamilton *et al*. 2015). However, the null *msl8-4* allele is in the Landsberg *erecta* (L*er*) background, while most mutants available from publicly available stock centers are in the Columbia-0 (Col-0) background. Furthermore, *MSL8* (At2g17010) has a close homolog, *MSL7* (At2g17000), located in tandem on Chromosome 2 (SI Appendix, Fig. S1A). MSL7 and MSL8 have partially overlapping expression pattern, as *MSL8* is expressed specifically in mature pollen gains and pollen tubes (Hamilton *et al*. 2015) and an *MSL7p::GUS* reporter was expressed in pistil stigma cells and pollen tubes when grown through the stigma and style (SI Appendix, Fig. S1B-C), consistent with microarray data (Swanson, Clark & Preuss 2005; Qin *et al*. 2009). We thus identified *msl7-1* (SALK_133223) as a null T-DNA insertion allele in the Col-0 background (SI Appendix, Fig. S1A, G-H). We then used CRISPR/Cas9 gene editing of the *MSL*8 gene to generate two null *msl8* alleles (*msl8-5* and *msl8-8)* and two double *msl7 msl8* mutants (*msl7-1 msl8-6* and *msl7-1 msl8-7)* in the Col-0 background (SI Appendix, Fig. S1D-F, Table S1). We also used artificial microRNA (amiRNA)-based silencing to reduce *MSL8* transcripts in the *msl7-1* mutant background. Both *msl7-1 amiR-MSL8-1* and *msl7-1 amiR-MSL8-2* showed reduced *MSL8* transcript levels in flowers (SI Appendix, Fig. S1G-H, Table S2).

We next germinated wild type and mutant pollen grains *in vitro* and monitored their morphology after 4 and 7 hours (h). We observed increased pollen grain bursting (with the visible release of cytosol) at both timepoints, and increased germination rates at the earlier timepoint, in the single *msl8* and double *msl7 msl8* mutant lines when compared to pollen from WT or *msl7-1* single mutant plants (Fig. 1A). Similarly, pollen grains from *msl7-1 amiR-MSL8-1* and *msl7-1 amiR-MSL8-2* showed significantly more bursting and early germination compared to WT or *msl7-1* (SI Appendix, Fig. S2A). There was no detectable difference between *msl8* single and *msl7 msl8* double mutant pollen in bursting or germination at either time point. The small but statistically significant difference in germination rates between *msl7-1* and WT pollen shown in Fig. 1A was not reproducible in other experiments (SI Appendix, Fig. S2).

**Figure 1.**
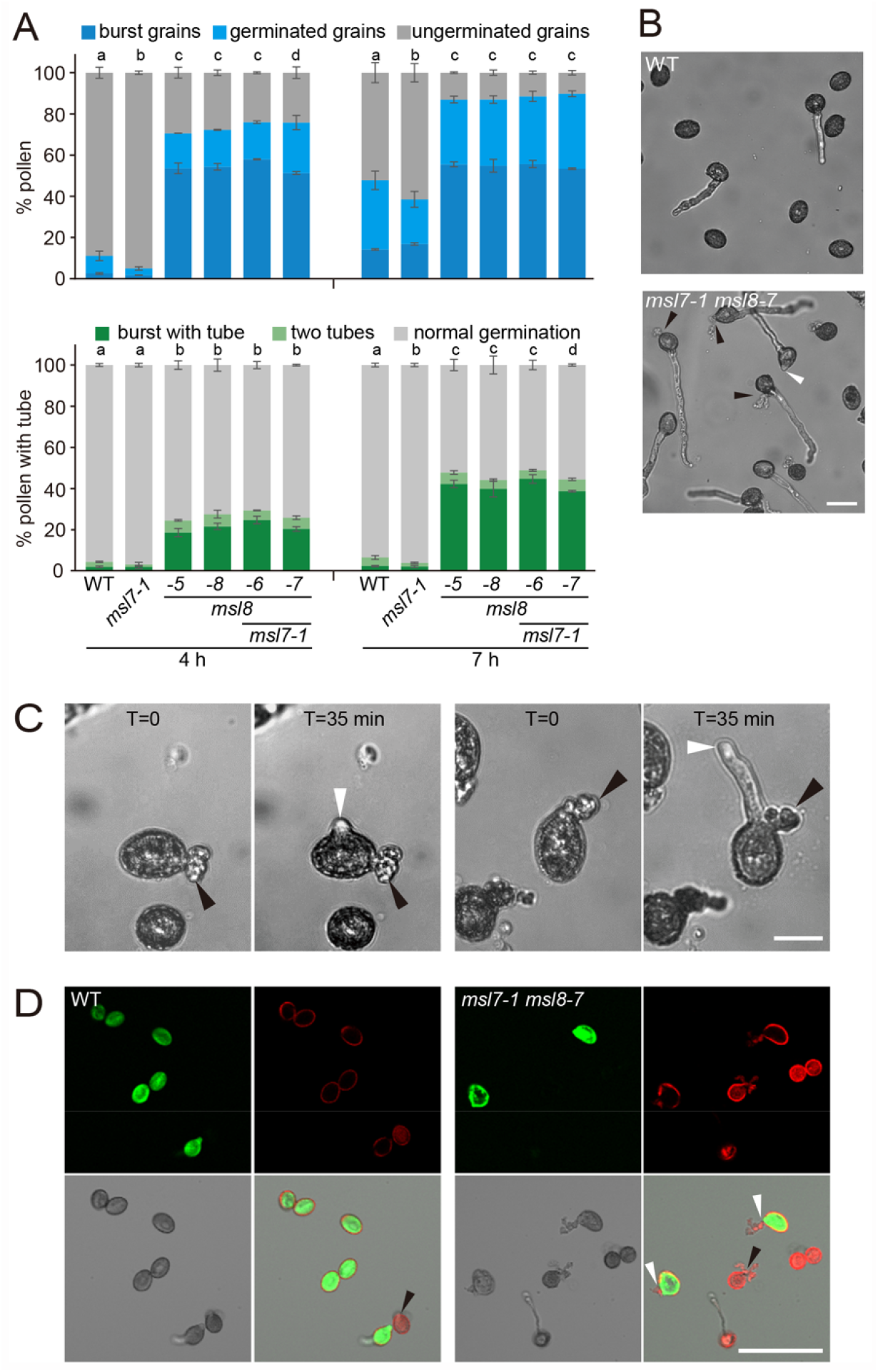
Pollen from *msl8* mutants show multiple phenotypes during *in vitro* germination. (A) Top: Bursting and germination rates at 4 and 7 h after *in vitro* germination in the indicated genotypes. Averages from 2 experiments with n = 1500-2000 grains per genotype per replicate are presented. Bottom: Abnormal germination phenotypes at 4 h and 7 h. Averages from 3 experiments with n = 100-300 grains per genotype per replicate at 4h and ∼500 grains at 7 h are presented. Error bars, standard error of the mean (SEM). Statistical analysis, Chi-square test, p < 0.001. (B) Brightfield images of pollen morphology after 4 h of *in vitro* germination. Black arrowheads, burst pollen grains with a tube. White arrowhead, two tubes from one pollen grain. Bar = 50 μm. (C) Time lapse images showing a burst pollen grain “resurrecting” to germinate a tube from *msl7-1 msl8-6* (left two panels) and *msl7-1 msl8-7* (right two panels) pollen. Black arrowheads, sites of pollen grain bursting; white arrowheads, a pollen tube growing from a burst pollen grain. Bar = 20 μm. (D) Confocal images of pollen after 2-3 h of *in vitro* germination in the indicated genotypes followed by viability staining. Green, Fluorescein diacetate (FDA); red, propidium iodide (PI). White and black arrowheads indicate alive and dead burst pollen grains, respectively. Bar = 50 μm.

### Several novel pollen germination phenotypes are associated with *msl7 msl8* mutants

A closer examination of germinated pollen grains from *msl8* and *msl7 msl8* CRISPR mutants revealed additional and unexpected phenotypes. Most WT germinated pollen grains were intact and produced a single pollen tube, while up to 50% of *msl8* and *msl7 msl8* mutant pollen grains either were burst but still had a growing tube or had germinated two tubes (Fig. 1A-B), which are collectively referred to here as “abnormal germination”. The same abnormal germination phenotypes were observed in pollen from both amiR-*MSL8* lines (SI Appendix, Fig. S2B). These phenotypes could be ecotype-dependent as they were not observed in *msl8-4* mutant in L*er* background.

To determine when bursting occurred, in the *msl8* and *msl7 msl8* pollen, we performed time lapse imaging of pollen grains in germination media. In pollen from the mutants, but not from the WT, we observed several cases of burst pollen grains that later “resurrected” an intact tube (Fig. 1C, Movies S1-3). We also observed examples of mutant grains with an intact tube producing a second growing tube (Movie S4). Thus, in some mutant grains, tubes could grow from grains that had burst before germination, and in some cases tubes could grow from grains that had already had a tube. To further explore this phenotype, we performed viability staining with fluorescein diacetate (FDA) for cell vitality and propidium iodide (PI) for membrane integrity (Boulos, Prévost, Barbeau, Coallier & Desjardins 1999; Muhlemann, Younts & Muday 2018). We found that 48% of burst *msl8* or *msl7 msl8* mutant pollen grains were still alive, compared to 23% of burst WT pollen (Fig. 1D). For all *msl8* mutant lines, the percentage of pollen that both burst and have a growing tube increased over time (Fig. 1A, SI Appendix, Fig. S2B). These observations suggest that grains lacking MSL8 have altered membrane and/or cell wall properties relating to turgor restoration and tube growth after grain bursting.

### MSL8 is required for normal intine structure and callose deposition at the germination site

To investigate a possible novel role for MSL8 in controlling cell wall properties, we used transmission electron microscopy to examine the ultrastructure of germinating WT and *msl7-1 msl8-7* pollen grains (i.e. with visible protrusions ranging from a small bump to a fully extended tube). We did not see obvious differences between the two genotypes with respect to pollen coat or exine but did observe striking changes in the intine at the germination site (Fig 2A). WT pollen had a relatively regular intine layer that thickened evenly at the germination site as reported previously (Hoedemaekers *et al*. 2015). In *msl7-1 msl8-7* pollen, the intine was dramatically disrupted; in some cases, it was abnormally thick (two headed arrows), while in other cases it looked discontinuous or had inclusions (black arrowheads). The inclusion could be cytoplasm as previously reported (Hoedemaekers *et al*. 2015). All these phenotypes indicate a role for MSL8 in establishing cell wall structure at the germination site.

**Figure 2.**
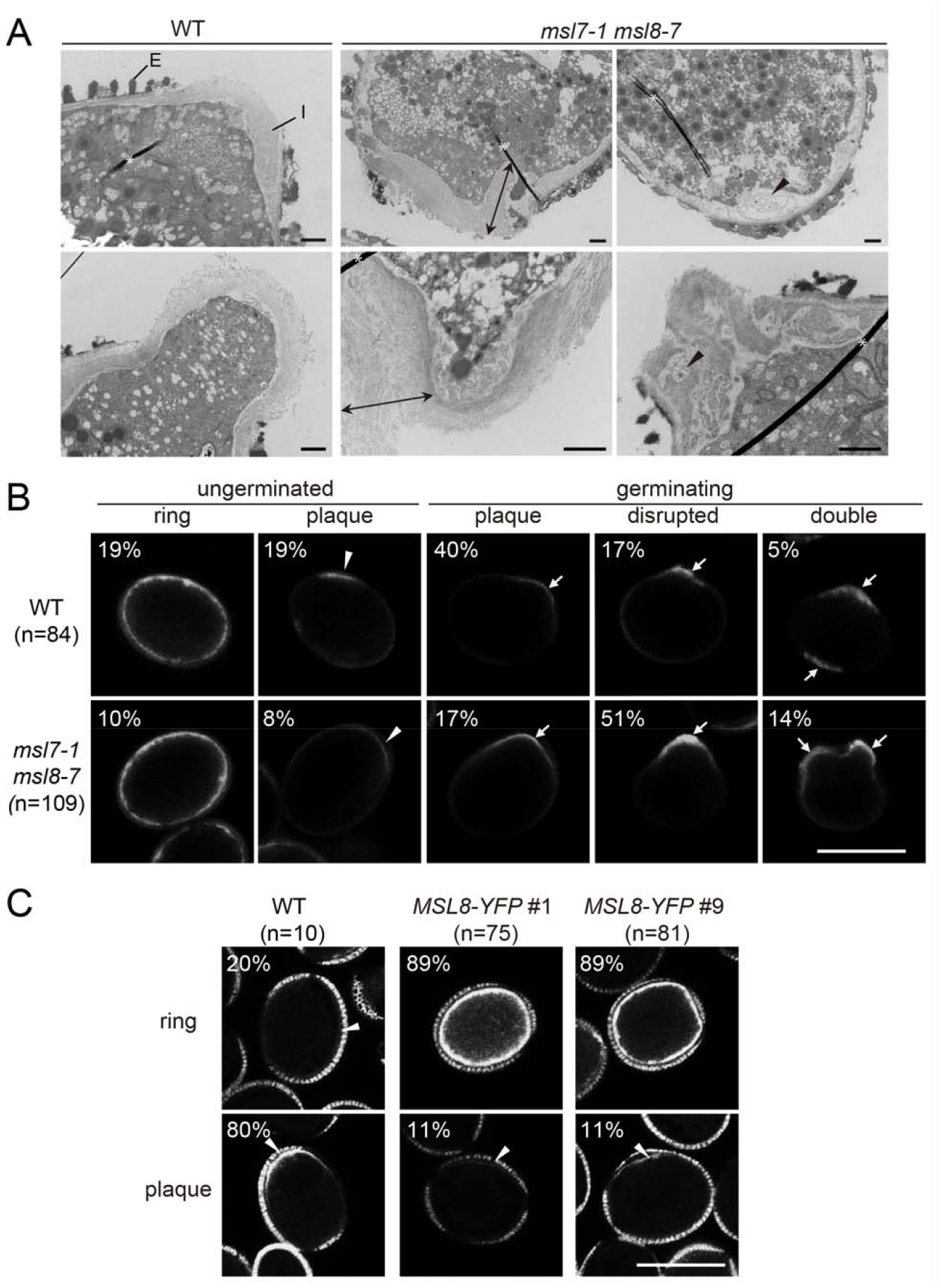
MSL8 is required for normal formation of the germination plaque. (A) Transmission electron microscopy images of germinating pollen grains from the indicated genotypes. In some cases, *msl7-1 msl8-7* grains exhibited extremely thick intine layers (double headed arrows) and in others disrupted intine with occasional inclusions (black arrowheads). Asterisk, section artifact. E, exine; I, intine. Scale bar = 1 μm. (B, C) Callose deposition in ungerminated or germinating pollen grains revealed by aniline blue staining. Due to very low germination rate in both MSL8-YFP lines, ungerminated and germinating pollen grains are combined. Germination sites, white arrows; potential germination sites, white arrowheads. Scale bars = 20 μm.

We next used aniline blue staining to visualize callose deposition in germinating and ungerminated WT and *msl7 msl8* mutant pollen grains after 2-4 h incubation in germination media (Fig. 2B). In both genotypes, many (mostly ungerminated) pollen grains remained unstained (81% in WT and 67% in *msl7-1 msl8-7*, SI Appendix, Fig. S3A). A lack of signal in these experiments could reflect an inability to trigger callose deposition in preparation for germination (Johnson & McCormick 2001) or to differences in the ability to react with aniline blue. Among those grains that stained with aniline blue, 59% of WT pollen grains showed a smooth callose plaque localized to the germination site or presumptive germination site, while only 25% of stained pollen from *msl7 msl8* mutants did (see ungerminated and germinating panels in Fig. 2B). Instead, the majority of these mutant pollen grains showed disrupted callose deposition, showing an uneven, concentrated or broadly distributed pattern at one (51%) or two sites (14%). A small portion of both genotypes (10-19%) showed a faint ring of callose around the grain (Fig. 2B).

We saw a different pattern in lines overexpressing *MSL8-YFP* from the strong, pollen-specific LAT52 promoter (*MSL8-YFP* hereafter) (Fig. 2C). Consistent with previous observations (Hamilton *et al*. 2015), the majority of pollen grains from both *MSL8-YFP* overexpression lines were ungerminated even after 4 h incubation in pollen germination medium (PGM). However, 47-69% of these ungerminated grains had detectable callose, a much higher rate than WT (8%, SI Appendix, Fig. S3A). Furthermore, most MSL8-YFP pollen grains that did stain with aniline blue showed a strong ring of callose around the grain (89%, Fig. 2C). A small population of WT (20%) pollen grains showed a similar pattern (Fig. 2C). We saw similar callose patterns in the *msl8-4* mutant and *MSL8-YFP* lines in the L*er* background (SI Appendix, Fig. S4). These data show that MSL8 is involved in the formation of the germination plaque and suggest that defects in cell wall properties could contribute to the abnormal germination phenotypes shown in Fig. 1.

### Overexpression of *MRI*^*R240C*^ suppresses *msl8* mutant phenotypes

The discovery of defective pollen cell walls in the *msl7-1 msl8-7* mutant led us to hypothesize that the pollen CWI pathway might be disrupted in these lines. We predicted that increased grain bursting and other phenotypes of *msl8* and *msl7 msl8* mutant pollen might be suppressed if CWI signaling was enhanced. To test this, we overexpressed the gain-of-function *MRI* allele *MRI*^*R240C*^ in WT and *msl7-1 msl8-7* mutant plants. We identified three independent lines expressing yellow fluorescence protein (YFP)- or cyan fluorescence protein (CFP)-tagged *MRI*^*R240C*^ under the pollen-specific *LAT52* promoter (Twell, Yamaguchi & McCormick 1990) in the *msl7-1 msl8-7* background (*msl7 msl8 MRI*^*R240C*^*-FP)*. Both fluorescence imaging (SI Appendix, Fig. S5) and immunoblotting (Fig. 3A) indicated a range of expression levels; lowest in line 5, intermediate in line 4 and highest in line 14. All *msl7 msl8 MRI*^*R240C*^*-FP* lines were backcrossed to WT plants, and siblings harboring homozygous *MRI*^*R240C*^*-FP* in either the WT or the *msl7 msl8* background were isolated. As expected, in the WT background, intermediate and high expression of *MRI*^*R240C*^*-FP* resulted in low bursting and low germination rates (Fig. 3B). Also as expected, *msl7 msl8* mutant pollen had high bursting and germination rates. Both phenotypes were significantly suppressed by intermediate and high *MRI*^*R240C*^*-FP* overexpression at both 4 and 7 h of incubation in germination media. The low level of *MRI*^*R240C*^*-FP* in line 5 also suppressed bursting and germination at the early time point, though not to WT levels. It is unclear why the low levels of *MRI*^*R240C*^*-FP* in line 5 in the WT background had higher bursting rates than the WT. We further tested if the abnormal germination phenotypes observed in *msl7 msl8* were also suppressed by overexpression of *MRI*^*R240C*^*-FP*. Indeed, expression of *MRI*^*R240C*^*-FP* suppressed abnormal germination compared to *msl7 msl8* in all three lines (Fig. 3C), primarily through reducing the rate of burst grains with a tube (Fig. 3C).

**Figure 3.**
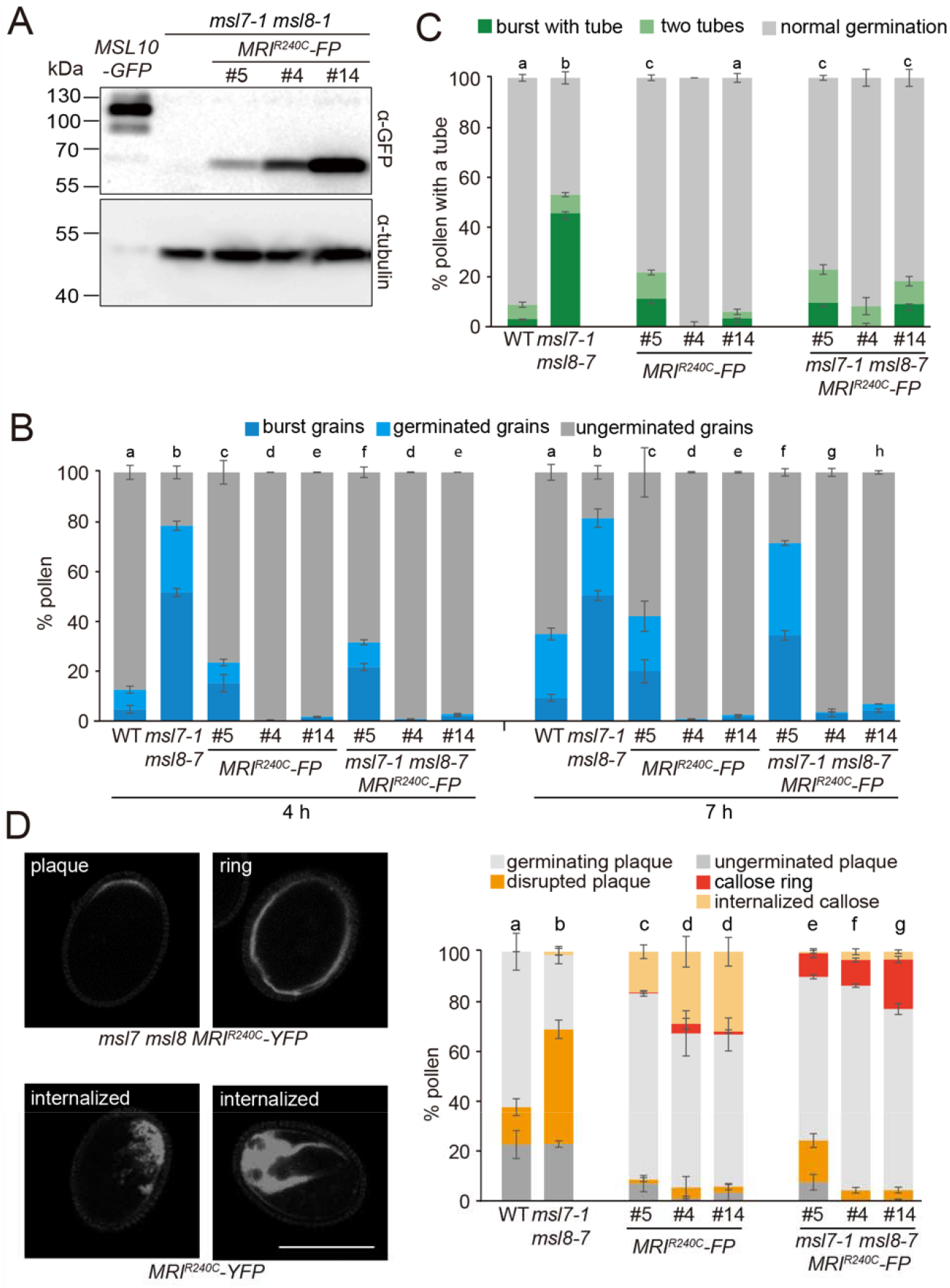
MARIS^R240C^ suppresses grain bursting and germination rates and alters callose deposition in *msl7 msl8* mutant pollen. (A) Immunoblot of total protein from flowers of the indicated genotypes. 35S::MSL10-GFP (Veley *et al*. 2014) was used as a positive control. The membrane was probed with anti-GFP antibody (upper panel) then stripped and re-probed with α-tubulin antibodies (bottom panel). (B) Pollen phenotypes during germination in the indicated genotypes. Averages from 3-6 experiments with n = 1000-2000 grains per genotype per replicate are presented. Error bars, SEM. (C) Abnormal pollen germination after 7 h in germination media. Averages from 3-6 experiments are present. N = 200-300 pollen tubes per replicate except for *MRI*^*R240C*^-FP #4 (n = 7), *MRI*^*R240C*^-FP #14 (n = 81), *msl7-1 msl8-7 MRI*^*R240C*^-FP #4 (n = 20), and *msl7-1 msl8-7 MRI*^*R240C*^-FP #14 (n = 175) due to low germination rates. Error bars, SEM. (D) Left, representative aniline blue-stained pollen grains from the indicated genotypes. Bar = 20 μm. Right, callose deposition pattern in the indicated genotypes. Only pollen grains with visible aniline blue stain are reported. Averages from 3-6 experiments with n = 50-100 grains per genotype per replicate are presented. Error bars, SEM. All *MRI*^*R240C*^ lines used in (A) were homozygous parental lines and in (B-D) were homozygous progenies isolated from the crosses between WT and *msl7-1 msl8-7 MRI*^*R240C*^. Statistical analysis in (B-D): Chi-square test, p < 0.05. Because of low sample numbers, statistical analysis was not carried out for *MRI*^*R240C*^-FP #4 nor *msl7-1 msl8-7 MRI*^*R240C*^-FP #s4 in (C).

The effect of *MRI* on grain bursting and germination in *msl8* prompted us to characterize its effect on callose deposition. The proportion of pollen grains that did not stain with aniline blue was reduced in the *msl7 msl8* background (SI Appendix, Fig. S3B). As shown above, almost half of the *msl7 msl8* mutant pollen that stained with aniline blue had a disrupted plaque pattern of callose staining (Fig. 3D). About 30% of pollen grains overexpressing *MRI*^*R240C*^*-FP* in WT background exhibited a dramatic pattern of internalized callose staining (Fig. 3D). However, when *MRI*^*R240C*^*-FP* was overexpressed in the *msl7 mls8* background, both the disrupted plaque phenotype of *msl7 msl8* and the internalized callose phenotype of *MRI*^*R240C*^*-FP* were absent, and most pollen grains either had a localized callose plaque or a ring around the periphery similar to the pattern seen in the MSL8-YFP overexpression lines (Fig. 3D). Internalized callose was rarely observed in the WT or *msl7 msl8* mutant lines. Taken together, the data shown in Fig. 3 indicate that the genetic relationship between *MSL8* and *MARIS* is not linear and are likely to reflect complex biochemical or biophysical pathways.

### Overexpression of *MSL8-YFP* suppresses grain bursting in *anx1/2* mutant pollen

To further test for genetic interactions between *MSL8* and the CWI pathway, we overexpressed *MSL8-YFP* in the *anx1/2* double mutant (*anx1/2 MSL8-YFP+/-*) and isolated 2 independent lines. The presence of YFP signal at the periphery and in internal puncta (as observed previously (Hamilton *et al*. 2015)) and immunoblotting indicated the presence of MSL8-YFP (Fig. 4A-B), but we were unable to isolate lines homozygous for the *MSL8-YFP* transgene, even after screening pollen from more than 50 T2 individuals. This may be due to suppressed male transmission efficiency in *MSL8-YFP* overexpression lines, as previously documented (Hamilton *et al*. 2015). We outcrossed both *anx1/2 MSL8-YFP+/-* lines to WT and identified siblings heterozygous for *MSL8-YFP* in both the WT and the homozygous double mutant *anx1/2* background.

**Figure 4.**
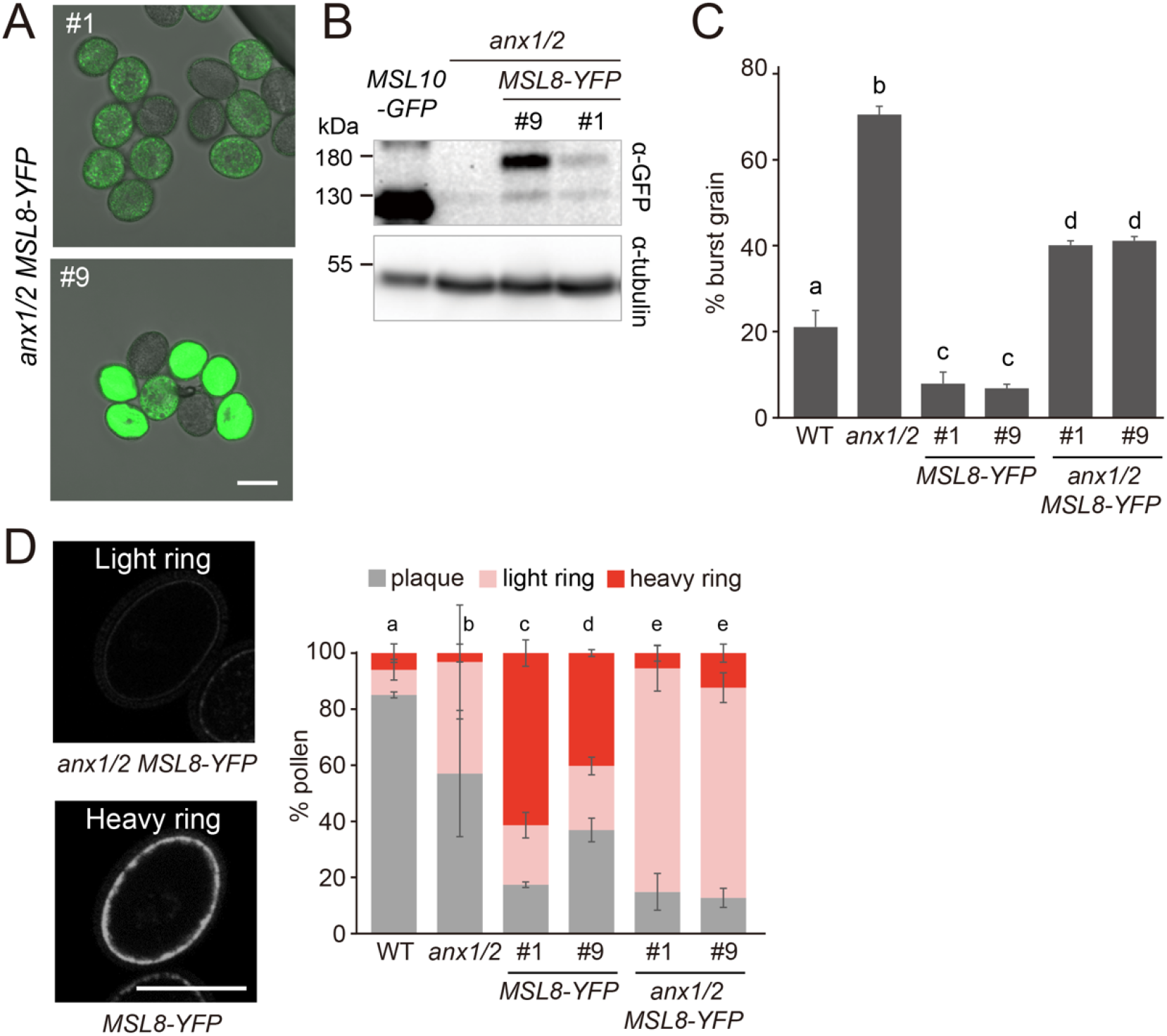
Overexpression of MSL8-YFP suppresses grain bursting and alters callose deposition in *anx1/2* mutant pollen grains. (A) Merged fluorescent and brightfield images showing YFP signal in pollen from two independent *anx1/2 MSL8-YFP+/-* lines. Bar = 20 μm. (B) Immunoblot of total protein from flowers of the indicated genotypes, performed as in Fig. 3A. (C) Pollen grain bursting rates at 24 h in the indicated genotypes. Averages from 3-7 experiments with n = ∼1000 pollen grains per genotype per replicate are presented. Error bars, SEM. Statistical analysis: ANOVA test, p < 0.05. (D) Left, representative images of callose deposition patterns in ungerminated pollen grains observed by aniline blue staining. Bar = 20 μm. Right, callose deposition pattern in the indicated genotypes. Only pollen grains with visible aniline blue stain are reported. Averages from 3 experiments with n = ∼30-120 per genotype per replicate are presented, except for *anx1/2* (n = 42 in total) due to high bursting rate. Error bars, SEM. Statistical analysis: chi-square test, p < 0.05. *MSL8-YFP* +/-lines were derived from a cross between WT and *anx1/2 MSL8-YFP* +/-.

We germinated parental and sibling pollen grains *in vitro* and examined their bursting rate (Fig. 4C). As expected, we observed a high degree of bursting in *anx1/2* double mutant pollen (∼70%) (Boisson-Dernier *et al*. 2009; Miyazaki *et al*. 2009), while both *MSL8-YFP+/-* lines showed a very low burst rate (∼8%) as previously described (Hamilton *et al*. 2015). In both *anx1/2 MSL8-YFP+/-*lines, we observed an intermediate amount of bursting. We speculated that this might be due to the segregation of the *MSL8-YFP* transgene in the pollen grains. Therefore, we used fluorescence microscopy to identify pollen with or without YFP signal and compared the pollen bursting rate in each background at different time points from 8 to 24 h. In pollen grains from *anx1/2 MSL8-YFP+/-* plants, the bursting rate in grains with YFP signal was always lower than those without a clear YFP signal (SI Appendix, Fig. S6).

To further characterize genetic interactions between *MSL8* and *ANX1/2*, we examined callose deposition in pollen grains from these lines by aniline blue staining. Since there were almost no intact germinating pollen grains in *anx1/2* mutant pollen, we focused on ungerminated pollen grains. As observed previously, not all pollen grains showed callose staining (SI Appendix, Fig. S3C). Additionally, the *anx1/2* alleles seemed to suppress staining even in the *MSL8-YFP* background (SI Appendix, Fig. S3C). Among the grains with callose signal, ∼85% of WT and ∼55% of *anx1/2* pollen grains showed a localized callose plaque (Fig. 4D). As above, overexpression of *MSL8-YFP* led to a heavy ring of callose around the pollen grain periphery. In *anx1/2 MSL8-YFP+/-* pollen grains, this ring was much fainter, suggesting that *ANX1/2* is partially required to produce the heavy callose ring in response to *MSL8-YFP* overexpression. Taken together, these findings further establish genetic interactions between *MSL8* and the pollen CWI pathway and suggest that they function in concert to maintain cell integrity during pollen germination.

## DISCUSSION

Plant cell expansion relies on a finely tuned balance between intracellular osmotic pressure and extracellular resistance from the cell wall, and dysregulation on either side of this equation can result in faulty growth behavior. In tip-growing cells, such as germinating and growing pollen tubes, the dynamic regulation of ion fluxes and of cell wall mechanical properties have been well-studied, but mostly separately (Michard, Simon, Tavares, Wudick & Feijó 2017; Cameron & Geitmann 2018). The work reported here shows that these two key mechanical contributions are inter-related, providing a novel avenue of mechano-homeostatic feedback and growth regulation.

We have proposed that MSL8 functions as an osmotic valve (Basu & Haswell 2017). According to this model, MSL8 is required to relieve internal pressure during pollen hydration and germination; in its absence, grains germinate more quickly and burst more readily. In overexpression lines, excess MSL8 releases pressure too efficiently, preventing the build-up of turgor that is required to drive germination. This “osmotic safety valve” model for MSL8 function in pollen is complicated by the phenotypes and genetic interactions presented here. Consistent with previous work (Hamilton *et al*. 2015), *msl8* or *msl7 msl8* mutants exhibited higher germination and bursting rates compared to the WT (Fig. 1A), while *MSL8-YFP* overexpression resulted in very low germination rates (Fig. 4C). However, we documented some new germination phenotypes, including double tubes and burst grains growing new tubes (Fig. 1A-C, SI Appendix, Fig. S2). A null *MSL7* allele did not affect any of these phenotypes.

### Evidence for a new role for MS ion channels in cell wall organization

The fact that mutant pollen grains could burst and then recover and germinate a growing tube while WT grains that had burst rarely did so (Fig.1D, Movies S1-3) suggested that there is more to MSL8 function than simply osmoregulation. Indeed, Transmission electron microscopy and aniline blue staining showed that MSL8 is required for normal cell wall organization in ungerminated pollen grains and at the germination site (Fig. 2, SI Appendix, Fig. S4). Pollen from *msl8* mutants showed a disrupted and disorganized germination plaque and patchy callose deposition at the germination site (Fig. 2, SI Appendix, Fig. S4). Furthermore, the accumulation of callose in a ring at the periphery of ungerminated pollen was much more frequently observed and much more dramatic in pollen from *MSL8-YFP* overexpressing plants than in pollen from the WT (Fig. 2C, Fig. 4D, SI Appendix, Fig. S4). Thus, based on TEM and if the presence of callose is used as a proxy for cell wall components, MSL8 unexpectedly modulates local cell wall organization in pollen grains before and during germination.

### Genetic interactions between MSL8 and the CWI pathway are complex

MSL8’s role in pollen integrity and cell wall deposition was at least superficially similar to that of the cell wall integrity pathway (Boisson-Dernier *et al*. 2009, 2013, 2015; Miyazaki *et al*. 2009), prompting us to test for genetic interactions between *MSL8* and two such components, *MRI* and *ANX1/2*. The results of these studies (summarized in Figure 5A) do not easily conform to linear genetic pathways. The grain bursting and abnormal germination phenotype of *msl8* pollen was suppressed by overexpression of *MRI*^*R240C*^*-FP* (Fig. 3A-C), while the grain bursting of *anx1/2* pollen was rescued by *MSL8-YFP* overexpression (Fig. 4A-C). In the *anx1/2* background, *MSL8-YFP* overexpression produced less intensely staining callose rings than they did in the *ANX1/2* background (Fig. 4D), suggesting that *ANX1/2* are required for the full effects of *MSL8-YFP* overexpression and that *MSL8* functions upstream of *ANX1/2*. However, other data suggest that MSL8 acts downstream of or parallel to ANX1/2. For example, excess callose was deposited to the periphery whether or not *ANX1/2* were present in *MSL8-YFP* overexpressing pollen (Fig. 4D).

**Figure 5.**
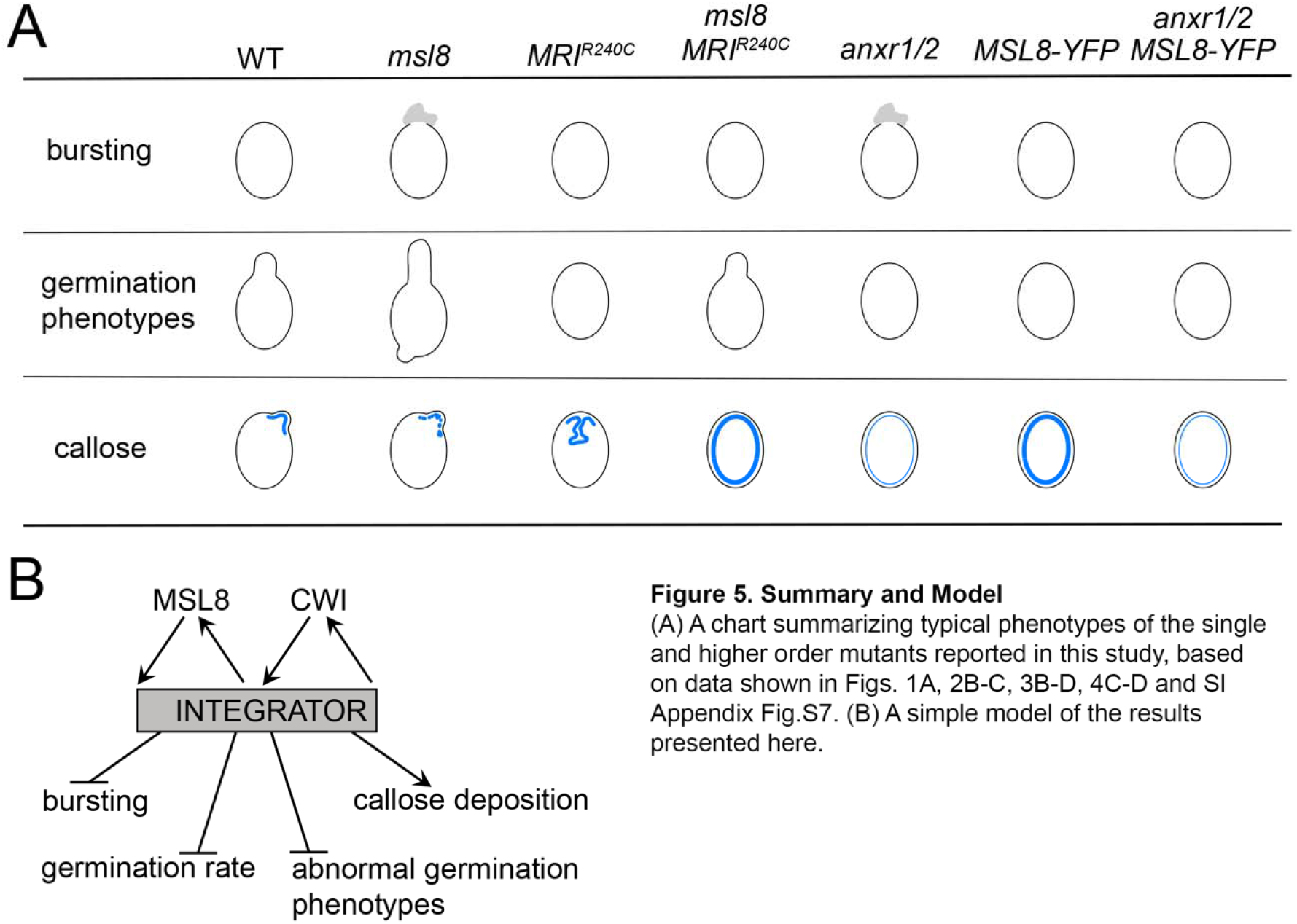
Summary and Model. (A) A chart summarizing typical phenotypes of the single and higher order mutants reported in this study, based on data shown in Figs. 1A, 2B-C, 3B-D, 4C-D and SI Appendix Fig.S7. (B) A simple model of the results presented here.

### Working model

Both MSL8 and the CWI pathway affect pollen germination, pollen bursting rate, and the pattern of callose deposition, and these phenotypes were affected in complex patterns that defy any linear relationship (Fig. 5A). As a result, we propose a simple model (Fig. 5B) wherein the MSL8 and the CWI signaling pathways are independent but impinge on a central common component, referred to here as an “integrator”, which can also feedback to either of them to ensure normal pollen germination and tube growth (Fig. 5B). This component could be a mechanical status, such as turgor, or a molecular signal, such as ions or reactive oxygen species, or a post-translational event such as the formation of a complex. Future work is needed to determine the exact mechanism by which MSL8 and the cell wall integrity pathway interact and how their signals are translated into downstream effects.

While it has become clear over the past few decades that mechanical forces play a huge role in plant development, how they play out at the cellular scale is not well understood. The results presented here indicate that in pollen, and likely in other plant cells, cell integrity maintenance systems require coordination between cell wall- and plasma membrane-based perception pathways and provide a foundation for future research into the mechanisms controlling mechanical homeostasis at the cellular scale.

## MATERIALS AND METHODS

### Plant growth

All the accessions used in this assay are Col-0 *Arabidopsis thaliana* unless otherwise specified. Surface sterilized seeds were plated on 0.5× Murashige and Skoog medium (MS) with appropriate antibiotics, incubated at 4°C for 2 days and then transferred to a long day chamber (Percival) at 21°C and 150 micromoles light (μmol) at 53% relative humidity for 6-7 days before transplanting to soil (Pro-Mix PGX Biofungicide, Premier Tech Horticulture). Plants were grown in long days at 21°C and 175 μmol at 45% relative humidity.

### Generation and identification of CRISPR mutants

Plasmid pHEE401E, with Cas9 driven by an egg cell-specific promoter (Wang *et al*. 2015), and pCBC-DT1T2 which facilitates the cloning of two guide RNAs (gRNA) (Xing *et al*. 2014) were used for CRISPR/Cas9 gene editing as described (Xing *et al*. 2014). Briefly, primers that harbored the two gRNA sequences targeting *MSL8* were used to amplify target fragments from pCBC-DT1T2, which was then introduced into pHEE401E yielding a destination construct for plant transformation. Two constructs, pHEE401E-MSL8_1e (LHP1198) and pHEE401E-MSL8_2e (LHP1199) (Table S1-2), were made using this method, and were introduced into Col-0 and *msl7-1* mutant plants, respectively, by floral dip for *Agrobacterium*-mediated transformation (Clough & Bent 1998). The transformants were screened on 0.5× MS plates supplemented with Hygromycin (35 mg/mL, Gold Biotechnology, H-270-1). Candidates were screened for insertion/deletions (indels) by polymerase chain reaction (PCR) amplification (Table S1) and gel electrophoresis, then confirmed by sequencing. Cas9 in the selected lines was removed by backcrossing and segregation in progeny was tested on antibiotic plates. Seedlings that showed Hygromycin-sensitive growth were transferred to 0.5× MS plates without hygromycin for recovery, then transplanted to soil. The absence of Cas9 was further confirmed by genotyping. Alternatively, Cas9 free individuals in the segregating progeny were directly isolated by genotyping. Primers used to detect Cas9 and other genotyping information for the new *msl8* and *msl7 msl8* CRISPR alleles can be found in Table S1.

### Generation and identification of amiRNA mutants

A *MSL8* amiRNA cassette from pAmiR-MSL8 (CSHL_030627) was recombined into a pollen-specific expression construct (pB7WGLAT52, a gift from Gregory Copenhaver) using Gateway technology to make LAT52::amiRNA MSL8. This construct was then introduced into the *msl7-1* mutant background by floral dip for *Agrobacterium*-mediated transformation (Clough & Bent 1998). T1 lines were screened for Basta resistance. Candidates were further screened for silencing of *MSL8* gene expression by RT-PCR using RNA extracted from 15-30 newly open flowers. RNA extraction was performed using an RNeasy Micro Kit (Qiagen). The full sequence for amiR-MSL8 including attB sites is shown in Table S2.

### Generation and identification of LAT52::MRI^R240C^-FP and LAT52::MSL8-YFP plants

LAT52::MRI^R240C^-YFP and LAT52::MRI^R240C^-CFP (Boisson-Dernier *et al*. 2015) and LAT52::MSL8-YFP (Hamilton *et al*. 2015) were introduced into *msl7-1 msl8-7* mutant plants or *anx1-2 anx2-1* homozygous plants (Boisson-Dernier *et al*. 2009; Miyazaki *et al*. 2009), separately, by floral dip for *Agrobacterium*-mediated transformation (Clough & Bent 1998) and screened for Basta resistance. The primers for confirming the presence of either construct in plants are listed in Table S1.

### *In vitro* pollen germination and imaging

About 40 newly open flowers were collected into a 1.7 mL Eppendorf tube with 1 mL pollen germination medium (PGM, 0.49 mM H_3_BO_3_, 2 mM Ca(NO_3_)_2_, 2 mM CaCl_2_, 1 mM KCl, 1 mM MgSO_4_, and 18% (w/v) sucrose with pH 7 adjust with KOH (Bou Daher, Chebli & Geitmann 2009)) and vortexed for 1 min before centrifugation at 10,000g for 3 min at room temperature. Then the pollen pellet was rinsed with 20 µL PGM, resuspended in fresh 20 µL PGM and transferred to a polyethylenimine (PEI, Sigma-181978) pre-treated Glass Bottom Microwell Dishes (MatTek, P35G-1.5-14-C) and dried until the grains were almost but not fully dry (1-5 min). Then 200 µL PGM was gently added to the petridish and it was placed in a humid box and incubated at 23°C (incubator, VWR, 89511-416) for 4 or 7 h. For PEI pretreatment, the glass bottom of the dish was covered with 100 µL diluted PEI (PEI : ddH_2_O = 1 : 30) for about 15 min before rinsing with 200 µL ddH_2_O twice. Then 200 µL PGM was added for 15 min before rinsing again with ddH_2_O and drying. Pollen germination images or movies were taken by Olympus IX73 microscope. For movies, 40× images were taken every 3 sec for 5-45 min.

### Pollen staining and imaging

Pollen viability staining was performed as described (Hamilton *et al*. 2015) using pollen that was incubated in PGM 2-3 h at 30°C before staining. Aniline blue staining was performed according to (Lu 2011; Liu *et al*. 2016) using 0.01% decolorized aniline blue (DABS) (Sigma-Aldrich) solution with slight modifications. Pollen was germinated in a 1.7 mL Eppendorf tube with 200 µL PGM for 4 h (Col samples) or 8 h (Ler samples) at 23°C before staining. Samples were stained in DABS for 4-6 h in dark before imaging. Confocal images were collected on Olympus FV3000 with the DAPI setting (excitation 405 nm and detection 430-470 nm).

### Transmission electron microscopy (TEM)

Pollen was collected and germinated in a 1.7 mL Eppendorf tube with 200 µL PGM at 23°C for 3.5 h (*msl7-1 msl8-7*) or 4.5 h (WT). Then pollen fixation and TEM was performed as described using chemical fixation method for pollen grains (Jia, Li & Yang 2017). After fixation, the following processes including infiltration, embedding, polymerization, ultrathin sectioning, staining and TEM imaging were processed in the Center for Cellular Imaging (WUCCI) at Washington University in St. Louis. Grids with sections were examined using JEOL JEM-1400 120kV TEM.

### Immunoblot analysis

Freshly opened flowers of tested genotypes and negative control were collected in a liquid nitrogen (LN2) pre-treated Eppendorf tube and frozen immediately in liquid nitrogen. 10-day old seedlings of 35S::MSL10-GFP transformants (Veley *et al*. 2014) were also collected as positive control. Protein preparation, immunoblot with an anti-GFP antibody (1:5000 dilution, Takara Bio) followed by and detection were performed as described (Basu, Shoots & Haswell 2020). Then the blot was stripped and re-probed with an anti-tubulin antibody (1:40,000 dilution, Sigma) for 0.5 h, followed by horseradish peroxidase (HRP)-conjugated goat anti-mouse secondary antibody (1:30,000 dilution; Millipore) incubation for 0.5 h. Antibody incubations were all performed at room temperature. Detection was performed using the SuperSignal West Dura Detection Kit (Thermo Fisher) and Syngene PXi 4 imager with GeneSys software version 1.4.1.0 (Syngene).

### Image analysis and processing

All image analysis and processing were performed in ImageJ (Fiji), PowerPoint, Adobe Photoshop CS6 or Adobe Illustrator 2019.

### Statistical analysis

Chi-square test was carried out for all categorical data sets and ANOVA for the others using R (R Core Team 2020) (https://www.r-project.org/). Letters represent statistical differences adjusted for multiple comparisons using the False Discovery Rate (FDR) method for chi-square test and Holm method for ANOVA test.

## ACKNOWLEDGEMENTS

We thank Dr. Qi-jun Chen (China Agricultural University, China) for pHEE401E and pCBC-DT1T2; Dr. Aurelien Boisson-Dernier (University of Cologne, Germany) for MRI^R240C^ constructs; Dr. Hannes Vogler (University of Zurich, Switzerland) for *anx1-2 anx2-1* seeds, Dr. Gregory Copenhaver (University of North Carolina at Chapel Hill, USA) for pB7WGLAT52, and the Center for Cellular Imaging at Washington University in St. Louis for TEM preparation, processing, and imaging. This research was supported by NSF MCB-1253103 and MCB 1929355 (to E.S.H), NSF DGE-1745038 (to K.M.) and the Center for Engineering Mechanobiology.

## AUTHOR CONTRIBUTIONS

YW and EH designed the experiments; YW, JC, KM and GJ performed the experiments; YW, JC and KM did the data analysis; YW, EH, JC and KM wrote the manuscript.

## DATA AVAILABILITY

The data that support the findings of this study are available from the corresponding author upon reasonable request.

## REFERENCES

Ackermann F. & Stanislas T. (2020) The plasma membrane–an integrating compartment for mechano-signaling. Plants 9.

Adhikari P.B., Liu X. & Kasahara R.D. (2020) Mechanics of Pollen Tube Elongation: A Perspective. Frontiers in Plant Science 11, 1–13.

Basu D. & Haswell E.S. (2017) Plant Mechanosensitive Ion Channels: An Ocean of Possibilities. Current Opinion in Plant Biology 40, 43–48.

Basu D., Shoots J.M. & Haswell E.S. (2020) Interactions between the N-And C-termini of the mechanosensitive ion channel AtMSL10 are consistent with a three-step mechanism for activation. Journal of Experimental Botany 71, 4020–4032.

Bidhendi A.J. & Geitmann A. (2019) Methods to quantify primary plant cell wall mechanics. Journal of Experimental Botany 70, 3615–3648.

Boisson-Dernier A., Franck C.M., Lituiev D.S. & Grossniklaus U. (2015) Receptor-like cytoplasmic kinase MARIS functions downstream of *Cr*RLK1L-dependent signaling during tip growth. Proceedings of the National Academy of Sciences 112, 12211LP–12216.

Boisson-Dernier A., Lituiev D.S., Nestorova A., Franck C.M., Thirugnanarajah S. & Grossniklaus U. (2013) ANXUR Receptor-Like Kinases Coordinate Cell Wall Integrity with Growth at the Pollen Tube Tip Via NADPH Oxidases. PLoS Biology 11.

Boisson-Dernier A., Roy S., Kritsas K., Grobei M.A., Jaciubek M., Schroeder J.I. & Grossniklaus U. (2009) Disruption of the pollen-expressed FERONIA homologs ANXUR1 and ANXUR2 triggers pollen tube discharge. Development 136, 3279–3288.

Bou Daher F., Chebli Y. & Geitmann A. (2009) Optimization of conditions for germination of cold-stored Arabidopsis thaliana pollen. Plant Cell Reports 28, 347–357.

Boulos L., Prévost M., Barbeau B., Coallier J. & Desjardins R. (1999) LIVE/DEAD® BacLight™: application of a new rapid staining method for direct enumeration of viable and total bacteria in drinking water. Journal of Microbiological Methods 37, 77–86.

Cameron C. & Geitmann A. (2018) Cell mechanics of pollen tube growth. Current Opinion in Genetics and Development 51, 11–17.

Chen X.Y. & Kim J.Y. (2009) Callose synthesis in higher plants. Plant Signaling and Behavior 4, 489–492.

Clough S.J. & Bent A.F. (1998) Floral dip: A simplified method for Agrobacterium-mediated transformation of Arabidopsis thaliana. Plant Journal 16, 735–743.

Fabrice T.N., Vogler H., Draeger C., Munglani G., Gupta S., Herger A.G., … Ringli C. (2018) LRX proteins play a crucial role in pollen grain and pollen tube cell wall development. Plant Physiology 176, 1981–1992.

Firon N., Nepi M. & Pacini E. (2012) Water status and associated processes mark critical stages in pollen development and functioning. Annals of Botany 109, 1201–1213.

Ge Z., Bergonci T., Zhao Y., Zou Y., Du S., Liu M.-C., … Qu L. (2017) Arabidopsis pollen tube integrity and sperm release are regulated by RALF-mediated signaling. Science 358, 1596–1600.

Ge Z., Cheung A.Y. & Qu L.J. (2019) Pollen tube integrity regulation in flowering plants: insights from molecular assemblies on the pollen tube surface. New Phytologist 222, 687–693.

Gigli-Bisceglia N., Engelsdorf T. & Hamann T. (2019) Plant cell wall integrity maintenance in model plants and crop species-relevant cell wall components and underlying guiding principles. Cellular and Molecular Life Sciences.

Hamant O. & Haswell E.S. (2017) Life behind the wall: Sensing mechanical cues in plants. BMC Biology 15, 1–9.

Hamilton E.S. & Haswell E.S. (2017) The Tension-sensitive Ion Transport Activity of MSL8 is Critical for its Function in Pollen Hydration and Germination. Plant and Cell Physiology 58, 1222–1237.

Hamilton E.S., Jensen G.S., Maksaev G., Katims A., Sherp A.M. & Haswell E.S. (2015) Mechanosensitive channel MSL8 regulates osmotic forces during pollen hydration and germination. Science 350, 438–441.

Hoedemaekers K., Derksen J., Hoogstrate S.W., Wolters-Arts M., Oh S.A., Twell D., … Rieu I. (2015) BURSTING POLLEN is required to organize the pollen germination plaque and pollen tube tip in Arabidopsis thaliana. New Phytologist 206, 255–267.

Höfte H. & Voxeur A. (2017) Plant cell walls. Current Biology 27, R865–R870.

Jia P.-F., Li H.-J. & Yang W.-C. (2017) Transmission Electron Microscopy (TEM) to Study Histology of Pollen and Pollen Tubes. In Plant Germline Development. Methods in Molecular Biology. (ed S. A.), pp. 181–189. Humana Press, New York, NY, New York.

Johnson M.A., Harper J.F. & Palanivelu R. (2019) A Fruitful Journey: Pollen Tube Navigation from Germination to Fertilization. Annual Review of Plant Biology 70, 809–837.

Johnson S.A. & McCormick S. (2001) Pollen germinates precociously in the anthers of raring-to-go, an Arabidopsis gametophytic mutant. Plant Physiology 126, 685–695.

Li C., Wu H.M. & Cheung A.Y. (2016) FERONIA and her pals: Functions and mechanisms. Plant Physiology 171, 2379–2392.

Li H.-J. & Yang W.-C. (2018) Ligands Switch Model for Pollen-Tube Integrity and Burst. Trends in Plant Science 23, 369–372.

Liao H.Z., Zhu M.M., Cui H.H., Du X.Y., Tang Y., Chen L.Q., … Zhang X.Q. (2016) MARIS plays important roles in Arabidopsis pollen tube and root hair growth. Journal of Integrative Plant Biology 58, 927–940.

Liu X., Castro C., Wang Y., Noble J., Ponvert N., Bundy M., … Palanivelu R. (2016) The role of LORELEI in pollen tube reception at the interface of the synergid cell and pollen tube requires the modified eight-cysteine motif and the receptor-like kinase FERONIA. Plant Cell 28, 1035–1052.

Lu Y. (2011) Arabidopsis Pollen Tube Aniline Blue Staining. Bio-Protocol 1, 4–5.

Maksaev G., Shoots J.M., Ohri S. & Haswell E.S. (2018) Nonpolar residues in the presumptive pore-lining helix of mechanosensitive channel MSL10 influence channel behavior and establish a nonconducting function. Plant Direct 2, e00059.

Michard E., Simon A.A., Tavares B., Wudick M.M. & Feijó J.A. (2017) Signaling with ions: The keystone for apical cell growth and morphogenesis in pollen tubes. Plant Physiology 173, 91–111.

Miyazaki S., Murata T., Sakurai-Ozato N., Kubo M., Demura T., Fukuda H. & Hasebe M. (2009) ANXUR1 and 2, Sister Genes to FERONIA/SIRENE, Are Male Factors for Coordinated Fertilization. Current Biology 19, 1327–1331.

Monshausen G.B. & Haswell E.S. (2013) A force of nature: molecular mechanisms of mechanoperception in plants. Journal of experimental botany 64, 4663–4680.

Muhlemann J.K., Younts T.L.B. & Muday G.K. (2018) Flavonols control pollen tube growth and integrity by regulating ROS homeostasis during high-temperature stress. Proceedings of the National Academy of Sciences 115, E11188–E11197.

Naismith J.H. & Booth I.R. (2012) Bacterial Mechanosensitive Channels — MscS□: Evolution’s Solution to Creating Sensitivity in Function. Annual Review of Biophysics 41, 157–177.

Parre E. & Geitmann A. (2005) More than a leak sealant. The mechanical properties of callose in pollen tubes. Plant Physiology 137, 274–286.

Peyronnet R., Tran D., Girault T. & Frachisse J.M. (2014) Mechanosensitive channels: Feeling tension in a world under pressure. Frontiers in Plant Science 5, 1–14.

Qin Y., Leydon A.R., Manziello A., Pandey R., Mount D., Denic S., … Palanivelu R. (2009) Penetration of the stigma and style elicits a novel transcriptome in pollen tubes, pointing to genes critical for growth in a pistil. PLoS Genetics 5.

R Core Team (2020) R: A language and environment for statistical computing, R Foundation for Statistical Computing, Vienna, Austria.

Ranade S.S., Syeda R. & Patapoutian A. (2015) Mechanically activated ion channels. Neuron 87, 1162–1179.

Sede A.R., Borassi C., Wengier D.L., Mecchia M.A., Estevez J.M. & Muschietti J.P. (2018) Arabidopsis pollen extensins LRX are required for cell wall integrity during pollen tube growth. FEBS Letters 592, 233–243.

Swanson R., Clark T. & Preuss D. (2005) Expression profiling of Arabidopsis stigma tissue identifies stigma-specific genes. Sexual Plant Reproduction 18, 163–171.

rinh D.C., Alonso-Serra J., Asaoka M., Colin L., Cortes M., Malivert A., … Hamant O. (2021) How Mechanical Forces Shape Plant Organs. Current Biology 31, R143–R159.

well D., Yamaguchi J. & McCormick S. (1990) Pollen-specific gene expression in transgenic plants: Coordinate regulation of two different tomato gene promotors during microsporogenesis. Development 109, 705–713.

Veley K.M., Maksaev G., Frick E.M., January E., Kloepper S.C. & Haswell E.S. (2014) Arabidopsis MSL10 has a regulated cell death signaling activity that is separable from its mechanosensitive ion channel activity. The Plant cell 26, 3115–3131.

Vogler H., Santos-Fernandez G., Mecchia M.A. & Grossniklaus U. (2019) To preserve or to destroy, that is the question: the role of the cell wall integrity pathway in pollen tube growth. Current Opinion in Plant Biology 52, 131–139.

Wang Z.P., Xing H.L., Dong L., Zhang H.Y., Han C.Y., Wang X.C. & Chen Q.J. (2015) Egg cell-specific promoter-controlled CRISPR/Cas9 efficiently generates homozygous mutants for multiple target genes in Arabidopsis in a single generation. Genome Biology 16, 1–12.

Wilson M.E., Jensen G.S. & Haswell E.S. (2011) Two mechanosensitive channel homologs influence division ring placement in Arabidopsis chloroplasts. Plant Cell 23, 2939–2949.

Xing H.-L., Dong L., Wang Z.-P., Zhang H.-Y., Han C.-Y., Liu B., … Chen Q.-J. (2014) A CRISPR/Cas9 toolkit for multiplex genome editing in plants. BMC Biology 14.

Xu T., Zhang C., Zhou Q. & Yang Z.N. (2016) Pollen wall pattern in Arabidopsis. Science Bulletin 61, 832–837.

