## Supplementary Files for "Interactions between a mechanosensitive channel and cell wall integrity signaling influence pollen germination in *Arabidopsis thaliana*"

* Elizabeth S. Haswell

**This PDF file includes:**

Figures S1 to S7 and corresponding figure legends

Table S1 to S2

Materials and Methods for Supplementary Information

References for Supplementary Information

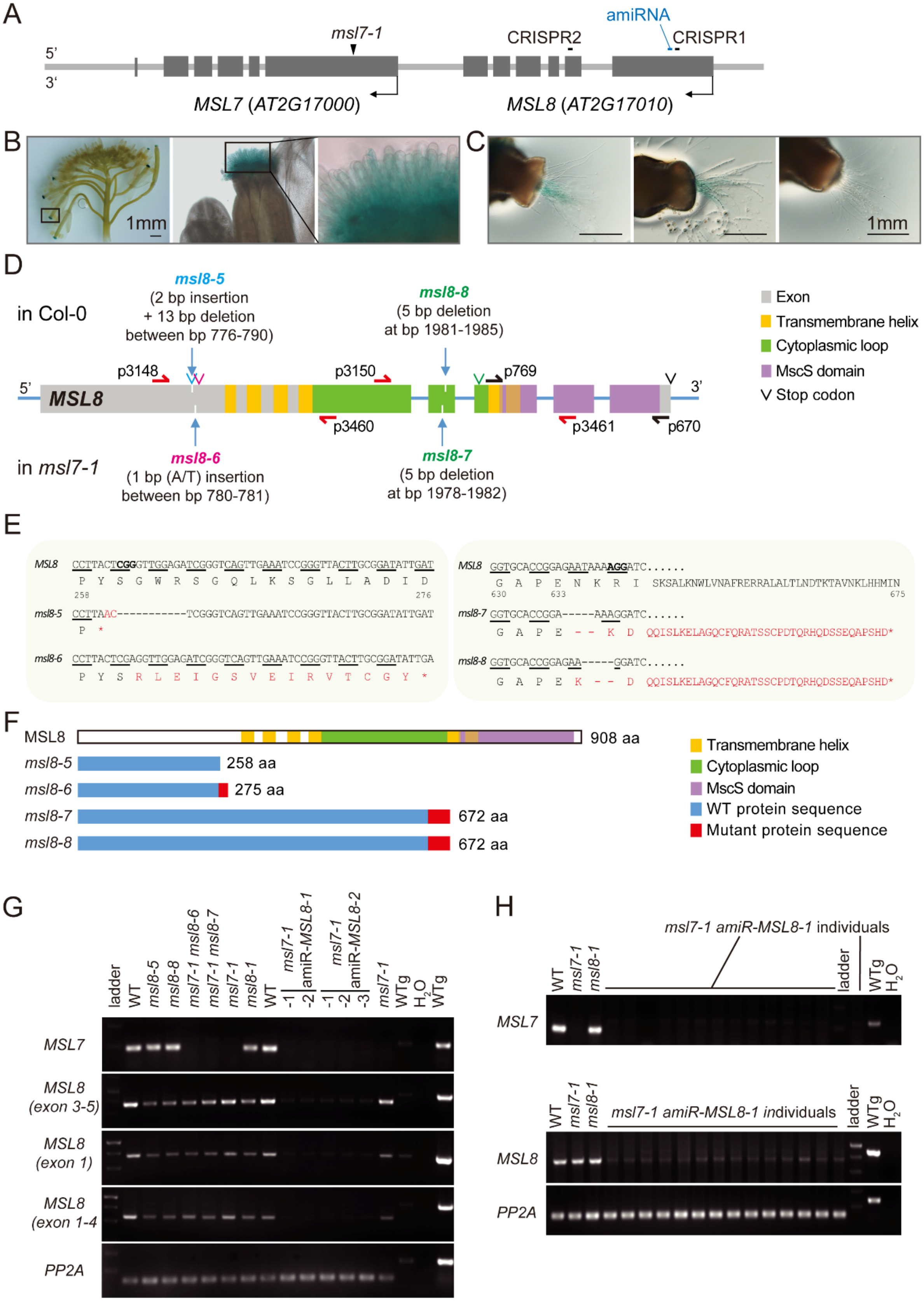
A
msl7-1
amiRNA
CRISPR2
CRISPR1
5'
3'
MSL7 (AT2G17000)
MSL8 (AT2G17010)
B
1mm
C
1mm
D
in Col-0
msl8-5
(2 bp insertion
+ 13 bp deletion
between bp 776-790)
msl8-8
(5 bp deletion
at bp 1981-1985)
p3148
p3150
p769
MSL8
p3460
p3461
p670
in msl7-1
msl8-6
(1 bp (A/T) insertion
between bp 780-781)
msl8-7
(5 bp deletion
at bp 1978-1982)
Exon
Transmembrane helix
Cytoplasmic loop
MscS domain
Stop codon
E
MSL8 CCTTACTCGGGTTGGAGATCGGGTCAGTTGAAATCCGGGTTACTTGCGGATATTGAT
P Y S G W R S G Q L K S G L L A D I D
258 276
msl8-5 CCTTAAC-----------TCGGGTCAGTTGAAATCCGGGTTACTTGCGGATATTGAT
P *
msl8-6 CCTTACTCGAGGTTGGAGATCGGGTCAGTTGAAATCCGGGTTACTTGCGGATATTGA
P Y S R L E I G S V E I R V T C G Y *
MSL8 GGTGCACCGGAGAATAAAAGGATC......
G A P E N K R I SKSALKNWLVNAFRERRALALTLNDTKTAVNKLHHMIN
630 633 675
msl8-7 GGTGCACCGGA-----AAAGGATC......
G A P E - - K D QQISLKELAGQCFQRATSSCPDTQRHQDSSEQAPSHD*
msl8-8 GGTGCACCGGAGAA-----GGATC......
G A P E K - - D QQISLKELAGQCFQRATSSCPDTQRHQDSSEQAPSHD*
F
MSL8 908 aa
msl8-5 258 aa
msl8-6 275 aa
msl8-7 672 aa
msl8-8 672 aa
Transmembrane helix
Cytoplasmic loop
MscS domain
WT protein sequence
Mutant protein sequence
G
ladder
WT
msl8-5
msl8-8
msl7-1 msl8-6
msl7-1 msl8-7
msl7-1
msl8-1
WT
msl7-1 amiR-MSL8-1 -1 -2
msl7-1 amiR-MSL8-2 -1 -2 -3
msl7-1
WTg
H2O
WTg
MSL7
MSL8 (exon 3-5)
MSL8 (exon 1)
MSL8 (exon 1-4)
PP2A
H
msl7-1 amiR-MSL8-1 individuals
WT
msl7-1
msl8-1
ladder
WTg
H2O
MSL7
MSL8
PP2A

**Figure S1. *MSL7* expression pattern and *msl8* mutants generated by CRISPR/Cas9 and amiRNA**. (**A**) A diagram illustrating the location of *MSL7* and *MSL8* in tandem on chromosome 2. Dark grey boxes, exons; light grey line, introns and non-coding sequences; arrows, transcription direction. CRISPR and amiRNA labels indicate the corresponding targeting sequences. (B-C) GUS-stained tissues from plants harboring *pMSL7:eGFP-GUS*. (**B**) Inflorescence. GUS blue signals appear in the stigmas of flowers from stage 12b-c (when the petals are longer than the stamens but flower bud is not open) to stage 16 (when petals and sepals wither) (Smyth, Bowman & Meyerowitz 1990; Christensen, King, Jordan & Drews 1997). The box in the left panel indicates the region shown in the middle panel; the box in the middle panel indicate the region shown in the right panel. GUS was not detected in any pollen grains. **C**) Pollen tubes growing through stigmas. GUS blue signals appear in pollen tubes when *pMSL7:eGFP-GUS* is present in the male parent. (**D**) Diagram of the *MSL8* gene and mutations generated for this study. Grey boxes, MSL8 coding sequence; yellow, transmembrane helix; green, cytoplasmic loop; purple, conserved MscS domain. Blue arrows, CRISPR/Cas9 genome editing sites; carets, stop codons; half arrows represent primer pairs used for RT-PCR in G and H. (**E**) DNA and corresponding protein sequence around the CRISPR/Cas9 gene editing sites of *MSL8*. PAM sequences are highlighted in bold. Red, mutant nucleotides or amino acids. Asterisk represents early stop codons. (**F**) Diagrams of predicted translation products from WT MSL8 and the four *msl8* CRISPR alleles. Red boxes indicate mutant sequences resulting from a change in reading frame. (**G, H**) Semi-quantitative RT-PCR analysis *MSL7* and *MSL8* transcripts in flowers from the indicated genotypes. g, genomic DNA. *msl8-1* is a partial loss-of-function allele (Hamilton *et al.* 2015).

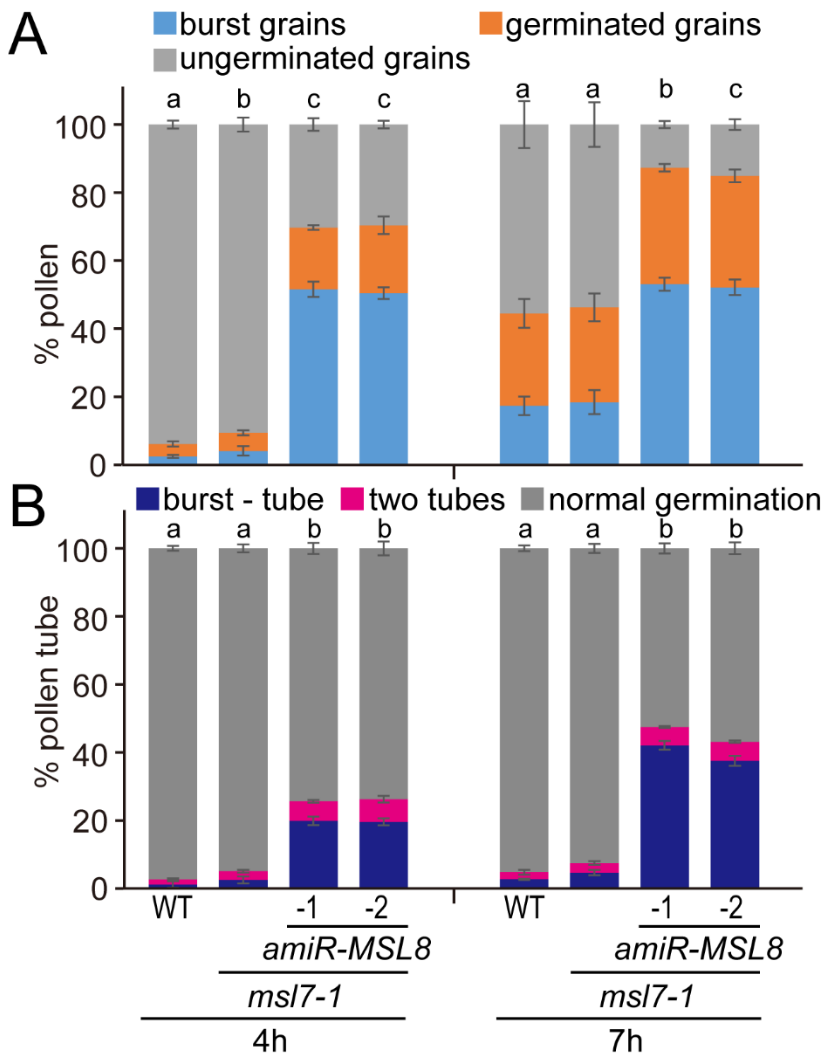

**Figure S2. Abnormal germination in *msl7 amiR-MSL8* lines.** (**A**) Pollen phenotypes during germination at 4 and 7 hours. Averages from 3 experiments with n= ~1500-2000 pollen per experiment are presented. (B) Abnormal germination at 4 and 7 hours. Averages from 3 experiments are presented. n= 100-300 pollen per genotype per experiment at 4 h and ~500 pollen per genotype per experiment at 7 h. Error bars, standard deviation (SD). Statistical analysis: chi-square test, p<0.001.

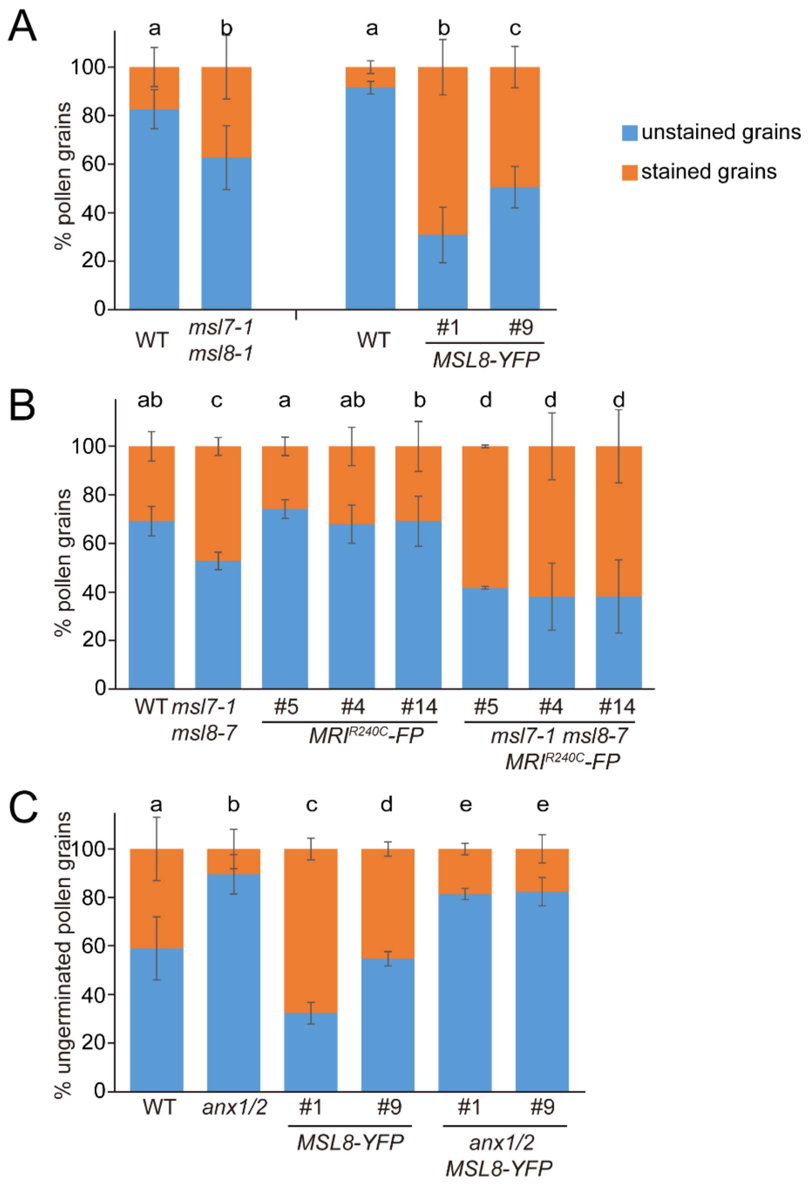

**Figure S3. Genotypic differences in pollen grain staining.** Pollen staining rate in *msl7 ms8* and *MSL8-YFP* experiments shown in Fig. 2B-C (**A**), in *MRI$^{R240C}$-PF* experiments shown in Fig. 3D (B), and in *MSL8-YFP* experiments shown in Fig. 4D (C). (A) Left, average from 3 experiments; Right, average from 3-8 imaging fields. n = ~ 150 pollen grains or more per genotype in total; (B) average from 3 experiments with n = ~150-200 per genotype per experiment. (C) average from 3 experiments with n = ~150 per genotype per experiment. Error bars, SD. Statistical analysis: chi-square test, $p<0.05$.

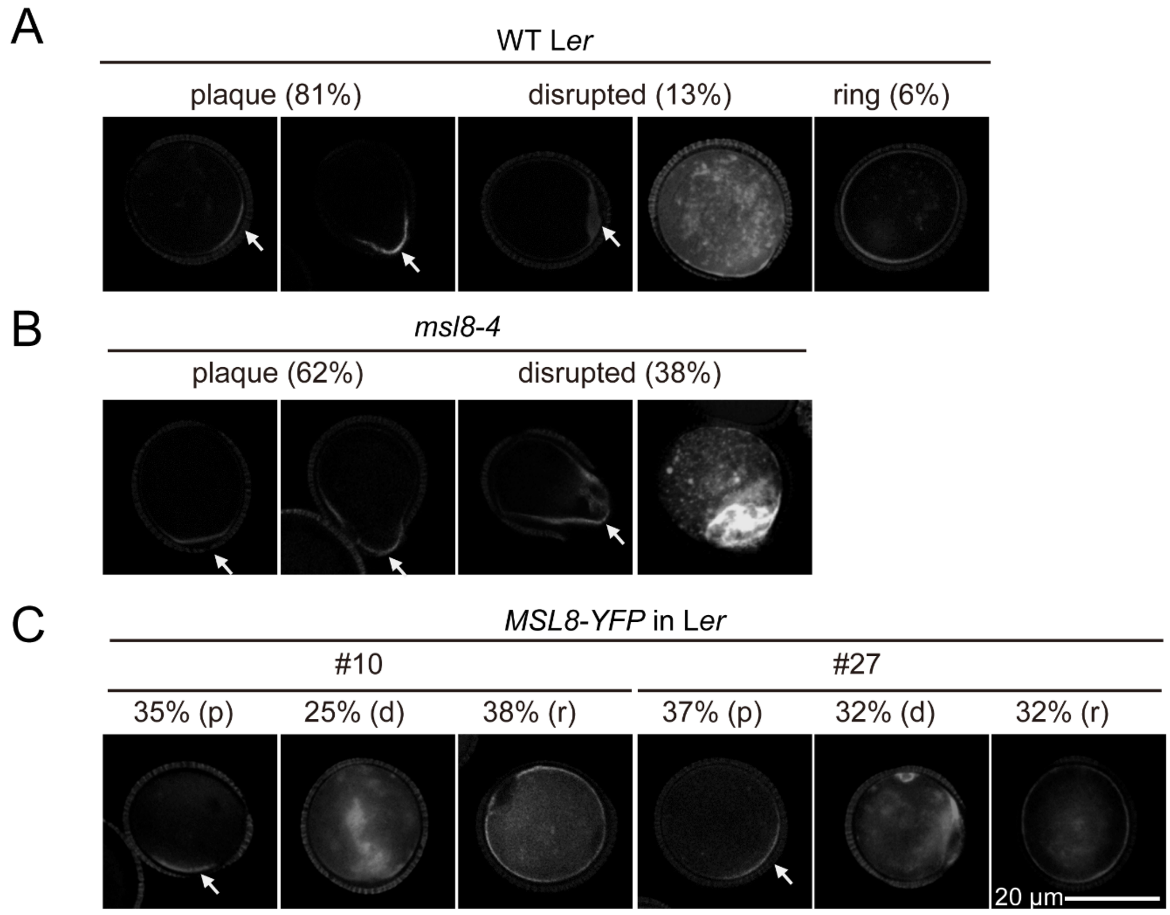

**Figure S4. Callose staining pattern in MSL8 loss-of-function or over-expression in the Landsberg *erecta* background.** Callose staining in pollen grains from WT (A, n = 67), *msl8-4* (B, n = 95), *LAT52:MSL8-YFP* line 10 (left three panels in C, n = 63) and *LAT52:MSL8-YFP* line 27 (right three panels in C, n = 57). Plaque (p), disrupted (d), ring (r).

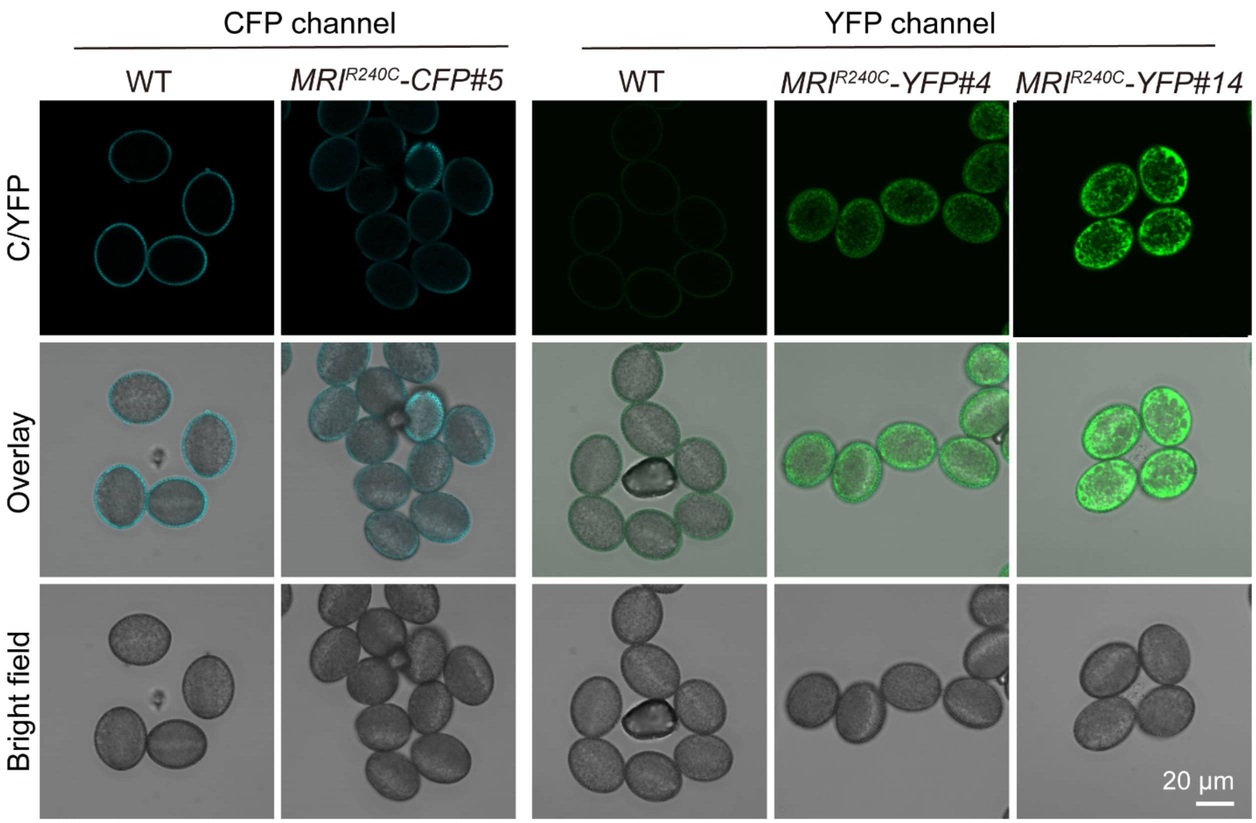

**Figure S5. Fluorescence imaging in *MRI$^{R240C}$-FP* lines.** Fluorescent signal in 3 independent lines of *msl8 MRI$^{R240C}$-FP*. Top panel, CFP or YFP fluorescence images (excited at 445 nm or 514 nm, respectively). Middle panel, merged fluorescence and bright field images. Bottom panel, bright field images.

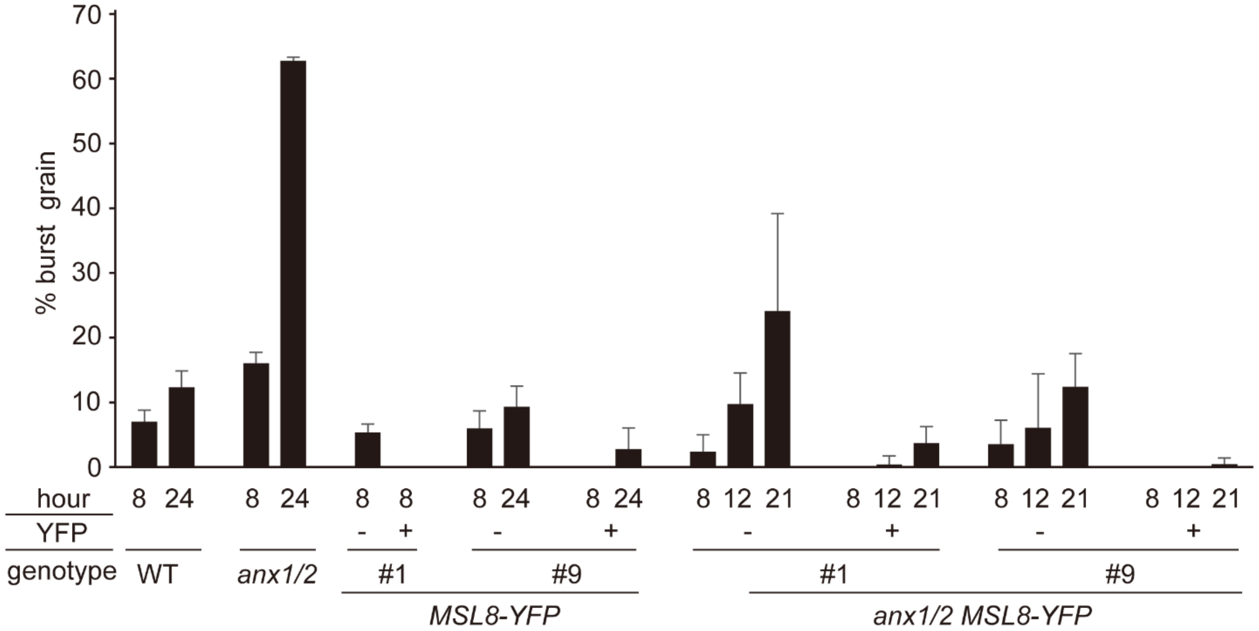

**Figure S6. MSL8-YFP overexpression suppresses pollen grain bursting in *anx1/2.*** Pollen grain bursting rate in the indicated genotypes at different time points after incubation (from 8 h to 24 h). Data from *MSL8-YFP* and *anx1/2 MSL8-YFP* were separated into grains with YFP signal (+) and grains without (-). Average from 4-14 imaging fields with n = ~100-200 pollen per genotype per field, except for *anx1/2 MSL8-YFP* (~50 pollen/field) Error bars, SD.

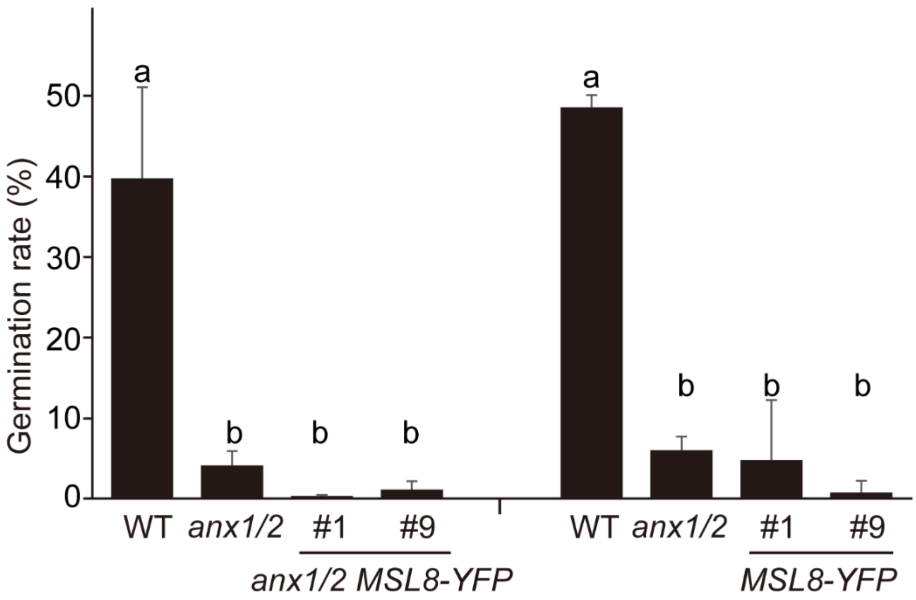

**Figure S7. Pollen germination rate in *anx1/2*, *MSL8-YFP* and *anx1/2 MSL8-YFP* backgrounds.** Left, average from 3-4 experiments with n = ~1000 pollen per genotype per experiment. Right, average from 4-9 imaging fields with n = 150-200 pollen per field for WT and *anx1/2*, and 150-200 pollen in total for *MSL8-YFP*. Error bars, SD. Statistical analysis, ANOVA, $p < 0.05$.

**Table S1**. Genotyping information.

| Allele | Locus | Primer set for gDNA | Primer set for T-DNA | Primer sequences | PCR | Gel % | Notes |
|---|---|---|---|---|---|---|---|
| *MSL8* | AT2G17010 | 1776+628 =422bp | | 1776: CGATCACCATCCTGATCATTCC<br>628: CTCTAGACGCATCAAACGAGGTTC<br>839: ACCCGACCGGATCGTATCGGT | 55℃ 20sec, 72℃ 40sec | 1% | |
| *msl8-4* DsLox N101568 | AT2G17010 | | 839 + 628 =~500bp | 1776: CGATCACCATCCTGATCATTCC<br>628: CTCTAGACGCATCAAACGAGGTTC<br>839: ACCCGACCGGATCGTATCGGT | 55℃ 20sec, 72℃ 40sec | 1% | |
| *MSL7* | AT2G17000 | 625+626 =511bp | | 625: ATGGAATTCCGCAAACCCTTTAAG<br>626: CCTTCGCCCTCGAAACTAGC<br>2229: ATTTTGCCGATTTCGGAAC | 55℃ 20sec, 72℃ 1min | 1% | |
| *msl7-1* SALK_133223 | AT2G17000 exon 1 | | 625 +1153/229 =~750bp | 625: ATGGAATTCCGCAAACCCTTTAAG<br>626: CCTTCGCCCTCGAAACTAGC<br>2229: ATTTTGCCGATTTCGGAAC | 55℃ 20sec, 72℃ 1min | 1% | |
| *msl8-6* [a] | AT2G17010 exon 1 (1A/T insertion) | 3402+3149 =95bp | | 3402: CACGTGAGGAAGAAACGCCTTACCCG<br>3149: CGCTAATGGATCATCCTCTTC | 58℃ 20sec, 72℃ 30sec | 3% | SmaI digestion (dCAPS) Only WT will be digested. |

| | | | | | | | |
|---|---|---|---|---|---|---|---|
| *msl8-7* [a] | AT2G17010 exon 2 (5bp deletion) | 3152+3153 =176bp [c] | | 3152: CCTCTGAATGACTAAAGGTAC 3153: TCTAATAAGACCAATGCAG | 50℃ 20sec, 72℃ 40sec | 3% | T7 Endonuclease I treatment (1 µl 15min) may be required to distinguish WT and homozygous |
| *msl8-5* | AT2G17010 exon 1 (13bp deletion + 2bp insertion) | 3148+3149 =167bp [d] | | 3148: CGAACATGTCGTTTCAGAGG 3149: CGCTAATGGATCATCCTCTTC | 55℃ 20sec, 72℃ 40sec | 3% | T7 Endonuclease I treatment (1 µl 20min) may be required to distinguish WT and homozygous |
| *msl8-8* | AT2G17010 exon 2 (5 bp deletion) | 3152+3153 =176bp | | 3152: CCTCTGAATGACTAAAGGTAC 3153: TCTAATAAGACCAATGCAG | 50℃ 20sec, 72℃ 40sec | 3% | T7 Endonuclease I treatment (0.5 µl 1h) may be required to distinguish WT and homozygous |
| LHP1198_pHEE401E-MSL8_1e;<br><br>LHP1199_pHEE401E-MSL8_2e [b] | Cas9 | 3248+3249 =~500bp | | 3248: GTACACGCGCAGGAAGAATC 3249: CAATGAGATTCCCGAACAGG | 55-56℃ 20sec, 72℃ 40sec | 1% | There may be non-specific band at ~400bp for Cas9-free individuals |
| MRI$^{R240C}$-FP [b] | AT2G41970 | | 3679+977 =684bp | 3679: GCATATGGAGCAGCCAAAGG 977: CTCGCCGGACACGCTGAACTTGT | 55℃ 20sec, 72℃ 60sec | 1% | For detection of MRI$^{R240C}$-FP insertion in the plant. |
| MSL8-YFP [b] | | | 3154+977= ~373bp or 768+977=~ 536bp | 3154: GAGTATTGGTACCCACAAGC 768: GCAATCGAGTTCTGTGTCCA | 55℃ 20sec, 72℃ 45sec | 1% | For detection of MSL8-YFP insertion in the plant. |

| | | | | |
|---|---|---|---|---|
| MSL7p::GFP-GUS [b] | | | 722+723=~3kb | 722: CACCATGAGTATAAAGAGGGAAGCTTG<br>723: GAAACAGAAACAGAAACTGTGAAGATGAT |

[a] these alleles are linked to *msl7-1*; we can genotype *msl7-1* to identify *msl8-5 and msl8-8*, as *MSL7* and *MSL8* are tandem in the genome.
[b] the names in the cells are for the constructs that the transgenic plants harbor.
[c] the primer set was also used for primary screening the indels in the first exon in *MSL8*.
[d] the primer set was also used for primary screening the indels in the second exon in *MSL8*.

**Table S2.** Sequence information for CRISPR and amiRNA constructs. Sequences in bold and underlined are those that target the MSL8 gene or transcript.

| | |
|---|---|
| Primer set 1 used to amplify gRNAs targeting exon 1 in *MSL8* | Forward primer 2880:<br>ATATATGGTCTCGATTG**AAGAAACGCCTTACTCGGGT**GTTTTAGAGCTAGAAATAGC<br>Reverse primer 2881:<br>ATTATTGGTCTCGAAAC**TTCGCTTCGTATCTGGCGGG**CAATCTCTTAGTCGACTCTAC<br><br>Note: Only the forward primer is functional. |
| Primer set 2 used to amplify gRNAs targeting exon 2 in *MSL8* | Forward primer 2882:<br>ATATATGGTCTCGATTG**AAGGTGCACCGGAGAATAAA**GTTTTAGAGCTAGAAATAGC<br>Reverse primer 2883:<br>ATTATTGGTCTCGAAAC**TGGTGATTAATCCTGTGACA**CAATCTCTTAGTCGACTCTAC<br><br>Note: Only the forward primer is functional. |
| Sequence sets used to target *MSL8* for artificial microRNA | Sequence 1 (amiR8*): **TAGCCACATTAGCGTGAAGTTTGTCGA**<br><br>Sequence 2 (amiR8): **TCGTCAAACTACACGCTAATGCGGCTA** |
| sequence for *amiR8* cassette (including attB sites) | ACAAGTTTGTACAAAAAAGCAGGCTCCACAAACACACGCTCGGACGCATATTACACATGTTC<br>ATACACTTAATACTCGCTGTTTTGAATTCATGTTTTAGGAATATATATGT**TAGCCACATTAGC**<br>**GTGAAGTTTGTCGA**CAGGTCGTGATATGATTCAATTAGCTTCCGACTCATTCATCCAAATAC<br>CGAGTCGCCAAAATTCAAACTAGACTCGTTAAATGAATGAATGATGCGGTAGACAAATTGGA<br>TCATTGATTCTCTT**TCGTCAAACTACACGCTAATGCGGCTA**CTCTTTTGTATTCCAATTTTCT |

| | |
|---|---|
| | TGATTAAGCTTTCCTGCACAAAAACATGCTTGATCCACTAAGTGACATATATGCTGCCTTCGT<br>ATATATAGTTCTGGTAAAATTAACATTTTGGGTTTATCTTTATTTAAGGCATCGCCATGGACC<br>CAGCTTTCTTGTACAAAGTGGT |

**Movies S1-3**.  Burst pollen grains from *msl7 msl8* mutant germinated a tube over time.

**Movie S4**. A grain with a tiny tube produce a different tube that is growing.

**Supplementary Methods**

**MSL7p::GUS reporter**. For MSL7p::GFP-GUS, the promoter fragment was amplified from 3 kilo base pair (kb) upstream of MSL7 using oligoes LHO722 and LHO723 (Table S1), cloned into pENTRY, sequenced, then recombined into pBGWFS7, trasnformed into plants, and selected with Basta. GUS staining was performed as described (Wang *et al.* 2017). Images were taken by stereoscope (Olympus SZX7) with DP71 camera and light microscope (Olympus, BX53) with DP80 camera.

**Semi-quantitative RT-PCR.** RNA was extracted by using either TRIzol Reagent (Invitrogen) or the RNeasy Mini RNA extraction kit (Qiagen) following the manufacturer's instructions. cDNAs were reverse transcribed from 1 µg RNA using an oligo$(dT)_{20}$ primer and M-MLV Reverse Transcriptase (Promega). For reverse transcriptase-polymerase chain reaction (RT-PCR), Taq polymerase and 1 µL of each cDNA were used in a 25 µL reaction. The corresponding primers for each target are listed in Table S1. 30 cycles of the following conditions were applied: 98 °C for 20 sec, and 55 °C for 20 sec, then 72 °C for 50 sec. At1g13320, which is a house-keeping gene encoding Protein Phosphatase 2A (PP2A) regulatory subunit, was used as an internal control (Czechowski, Stitt, Altmann, Udvardi & Scheible 2005). When running electrophoresis, 15 µL of the PCR products from each test sample was loaded, while 5 µL of control and genomic DNA were loaded.

**Supplementary References**

Christensen C.A., King E.J., Jordan J.R. & Drews G.N. (1997) Megagametogenesis in Arabidopsis wild type and the Gf mutant. *Sexual Plant Reproduction* **10**, 49–64.

Czechowski T., Stitt M., Altmann T., Udvardi M.K. & Scheible W.-R. (2005) Genome-Wide Identification and Testing of Superior Reference Genes for Transcript Normalization in Arabidopsis. *Plant Physiology* **139**, 5 LP – 17.

Hamilton E.S., Jensen G.S., Maksaev G., Katims A., Sherp A.M. & Haswell E.S. (2015) Mechanosensitive channel MSL8 regulates osmotic forces during pollen hydration and germination. *Science* **350**, 438–441.

Smyth D.R., Bowman J.L. & Meyerowitz E.M. (1990) Early flower development in Arabidopsis. *Plant Cell* **2**, 755–767.

Wang Y., Tsukamoto T., Noble J.A., Liu X., Mosher R.A. & Palanivelu R. (2017) Arabidopsis LORELEI, a maternally expressed imprinted gene, promotes early seed development. *Plant Physiology* **175**, 758–773.
